# Optimising digital volume correlation across materials: a practical framework for accuracy and spatial resolution

**DOI:** 10.64898/2026.09.15.751389

**Authors:** Alissa L. Parmenter, A. Sharma, B. K. Bay, P. D. Lee

**Affiliations:** Multiscale X-ray Imaging Lab, Department of Mechanical Engineering, University College London, Roberts Engineering Building, University College London, Torrington Place, London WC1E 7JE, UK; Department of Medical Physics and Bioengineering, Malet Place Engineering Building, University College London, Gower Street, London WC1E 6BT, UK; Research Complex at Harwell, Rutherford Appleton Laboratory, Harwell Oxford, Didcot OX11 0FA, UK; School of Mechanical, Industrial, and Manufacturing Engineering, 204 Rogers Hall 2000 SW, Monroe Avenue, Oregon State University, Corvallis, OR 97331-6001, USA

## Abstract

Digital volume correlation (DVC) provides full-field three-dimensional measurements of internal deformation, but its accuracy and effective spatial resolution depend on image-processing and analysis choices. Here, we use fibrous, cartilaginous and mineralised tissues within a rat intervertebral disc (IVD) as a controlled multi-material case study, combining virtual compression with experimentally loaded synchrotron computed tomography images. We systematically evaluate phase retrieval and image filtering, bit-depth conversion, point-cloud design, subvolume size and strain-field smoothing. Stronger phase retrieval reduced correlation residuals while increasing displacement and strain errors, showing that residual minimisation alone can select poorer parameters. Image filtering sensitivity was greatest in fibrous tissue, which had the smallest characteristic image feature size, whereas inappropriate intensity mapping during 16-bit to 8-bit conversion preferentially degraded low-contrast cartilage. Increasing subvolume size, point spacing or strain-window size improved measurement robustness but progressively smoothed local strain heterogeneity. We demonstrate that the spatial resolution of strain measurement must be matched across tissue types in order to compare strain magnitude; in the IVD, matching DVC spatial resolution changed the apparent ratio of compressive strain among fibrous, cartilaginous and mineralised tissues from 3.1:1.9:1 to 14:7.7:1. These findings establish a sequential, deformation-based optimisation framework in which image characteristics guide processing, known deformations validate accuracy and strain fields are compared at matched measurement scales. The framework supports more reproducible and mechanically interpretable DVC analyses in heterogeneous biological and engineered materials.

## Introduction

Understanding how deformation is distributed within complex multi-material systems is central to linking structure to mechanical function. In recent years, digital volume correlation (DVC) has emerged as a powerful approach for measuring full-field three-dimensional displacement and strain fields [1], with X-ray computed tomography the most commonly used imaging modality. DVC is now widely applied to investigate deformation mechanisms across a broad range of biological and engineering systems, including intervertebral disc [2–4], bone [5–9], cartilage [10, 11], lung [12], metallic [13] and composite [14] materials. By enabling non-destructive quantification of internal strains, DVC provides a unique opportunity to decipher the links between microstructural architecture and mechanical behaviour.

The accuracy and spatial resolution of DVC measurements depend strongly on user-selected processing parameters. Key choices include image filtering, sub-volume size, point-cloud density, and strain calculation methodology. These parameters control a complex trade-off between correlation robustness, measurement noise, computational cost and the spatial scale at which deformation can be resolved. Although numerous DVC studies report parameter optimisation for individual materials or experimental systems [11, 15], optimisation procedures are often based on empirical trial-and-error approaches, making it difficult to compare results between studies or establish reproducible best practices. Furthermore, commonly used optimisation metrics such as correlation residuals are frequently assumed to reflect measurement accuracy, despite limited evidence that this assumption remains valid across different image-processing conditions and material classes.

The challenge is particularly acute for biological materials, which span a wide range of microstructural organisations and imaging characteristics. Mineralised tissues contain high-contrast heterogeneous features that are generally favourable for image correlation, whereas soft tissues often exhibit lower contrast [16, 17]. Fibrous materials present an additional challenge as their highly anisotropic texture can generate elongated minima in the correlation landscape, increasing susceptibility to displacement tracking errors along fibre directions [18]. The mechanics of fibre-based materials is of interest at both the fibre and bulk material scales, and we have recently demonstrated that accurate measurements of both fibre and bulk-tissue level strain can be obtained by using a fibre-based DVC point cloud with different methods of strain calculation [2].

Parameter choices that are optimal for one tissue type may perform poorly in another, even when imaged under identical conditions. A general framework linking tissue microstructure, image characteristics and DVC performance across multi-material systems is therefore needed.

Here, we used three structurally distinct musculoskeletal tissues captured within the same synchrotron computed tomography (sCT) image volume as a controlled model system: collagen fibres of the annulus fibrosus (AF), hyaline cartilage of the cartilage endplate (CEP), and mineralised tissues of the vertebral endplate (VEP) (Fig. 1a). These tissues differ markedly in image contrast and characteristic feature size (Fig. 1b), allowing the influence of image structure on DVC performance to be evaluated independently of acquisition conditions. Using the open-source software iDVC, we systematically investigated how image filtering, point-cloud design and spatial-resolution parameters affect displacement and strain error, strain-field reproducibility and effective spatial resolution in virtual-deformation and experimental-loading analyses. By linking these outcomes to tissue-specific image characteristics, we identify the factors governing DVC performance and develop a practical framework for selecting parameters in complex, heterogeneous materials.

**Figure 1.**
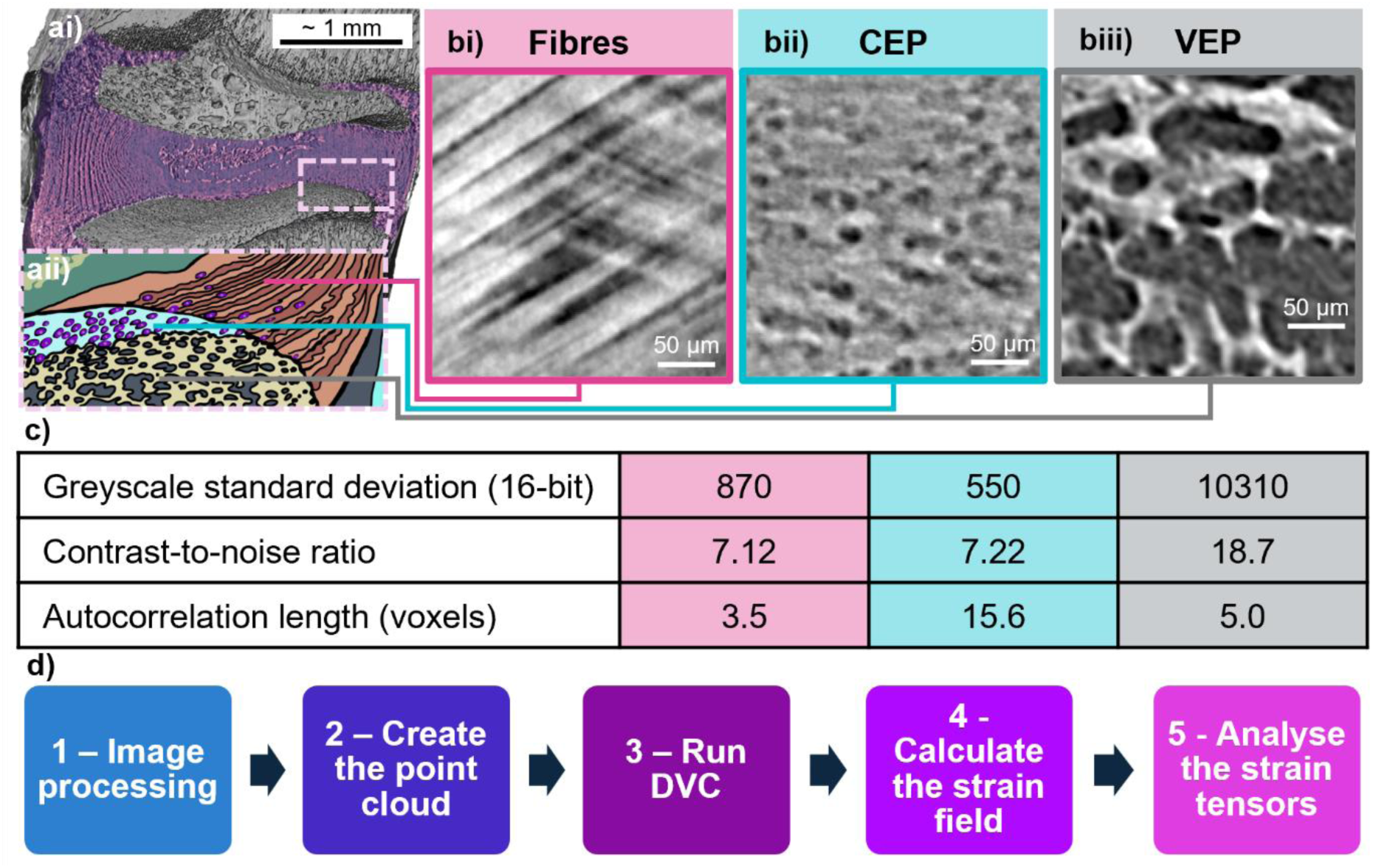
Microstructurally distinct tissues in the IVD display different image characteristics. ai) 3D rendering of the intervertebral disc with aii) illustration of the AF-CEP-VEP junction; b) greyscale sCT images of i) fibres in the annulus fibrosus, ii) hyaline cartilage in the cartilage endplate, and iii) mineralised tissues (bone and cartilage) in the vertebral endplate; c) image characteristics of the three tissue types; d) a flow chart showing the steps to performing DVC analysis on *in situ* images.

## Results and Discussion

Phase-contrast enhanced synchrotron computed tomography (sCT) images of rat intervertebral discs under compressive load were acquired at the I13-2 beamline, Diamond Light Source with a voxel size of 1.6 μm as described previously [8] (Methods). The experiment was designed to generate images with a high enough quality to accurately perform DVC in both hard and soft tissues whilst minimising radiation dose, with mechanical loading designed to be physiologically relevant whilst still being trackable by DVC [8]. This study focuses on optimising the DVC analysis pipeline after image acquisition; studies discussing optimisation of image acquisition are available elsewhere [16–20].

To establish how tissue-dependent image characteristics influence DVC performance, we first quantified the structural and greyscale characteristics of the AF, CEP and VEP. The mineralised VEP tissues (bone and mineralised cartilage) exhibited 12 to 18 times greater greyscale-intensity standard deviation and a 2.5-fold higher contrast-to-noise ratio than the soft tissues of the CEP and AF (Fig. 1c).

The characteristic feature scale, defined by the 1/e decay length of the radial autocorrelation function (Methods, Supplementary Fig. 1), was smallest in the fibrous AF tissue at 3.5 voxels (5.6 μm), consistent with the approximately 5 μm diameter of the collagen fibres (Fig. 1bi). By contrast, the CEP had the largest characteristic feature scale at 15.6 voxels (25.4 μm), reflecting the approximately 25 μm diameter and spacing of the chondrocytes (Fig. 1b(ii)). These tissues therefore spanned marked differences in both image contrast and structural length scale, enabling us to determine how each property constrains the accuracy and spatial resolution achievable by DVC. The following sections explore how parameter choices at each stage of the DVC analysis pipeline (Fig. 1d) differently impact the DVC measured deformation fields across these tissues.

### 1 Image processing

#### Phase retrieval alters DVC accuracy

After the experimental image acquisition phase, image processing is the first step in the DVC analysis pipeline which can have an impact on strain measurement accuracy. Phase retrieval algorithms are commonly applied in phase-contrast X-ray tomography to enhance image contrast and suppress noise, typically using the Paganin method [21]. This smooths out phase fringes, improving image contrast whilst suppressing noise but resulting in image blur (Fig. 2a). An unsharp mask is commonly applied to reduce image blur [8, 22].

**Figure 2.**
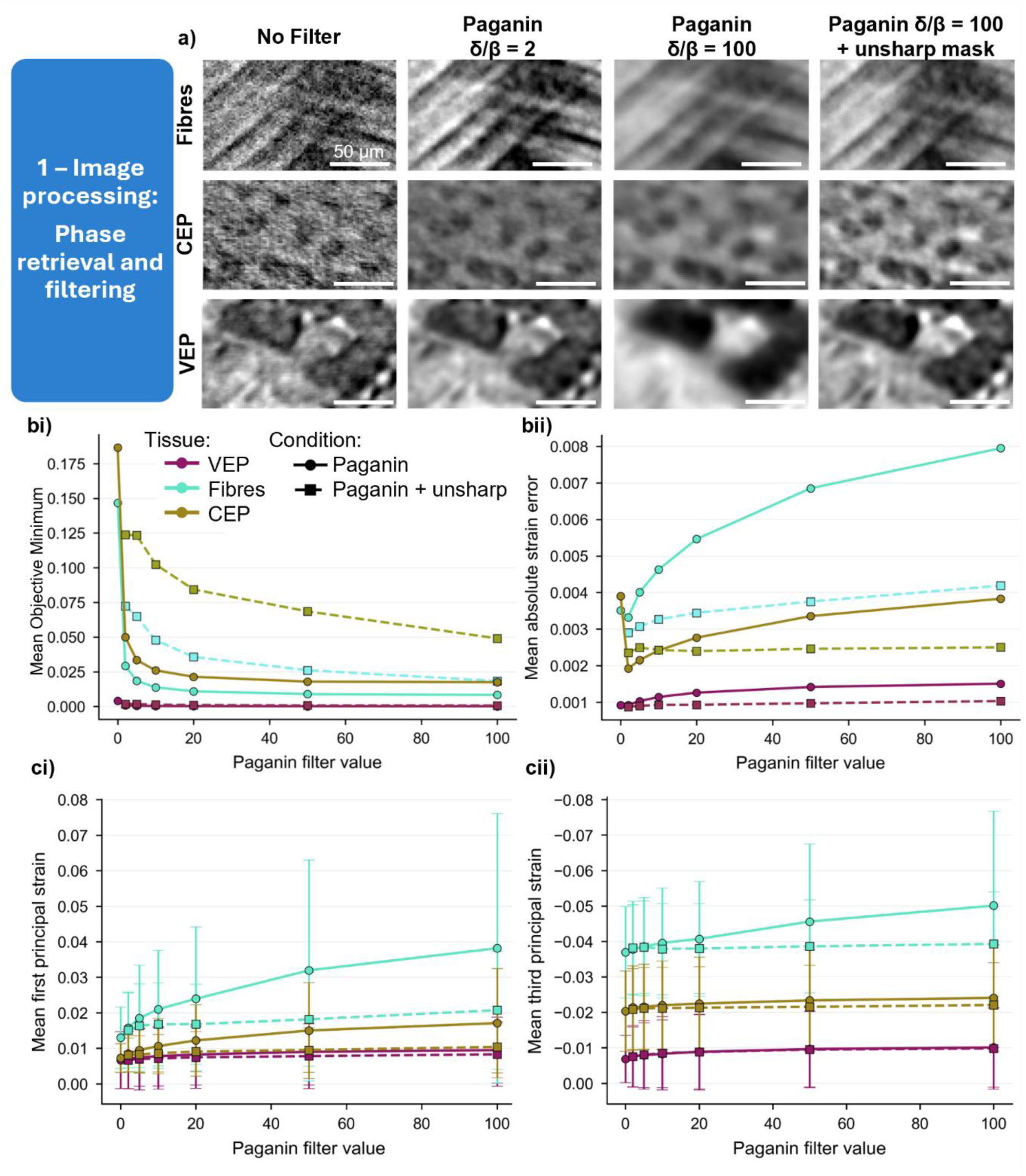
Image processing: The effect phase retrieval and filtering, a) the impact of Paganin phase retrieval and unsharp mask application on greyscale sCT images of VEP, AF fibres, and CEP; b) the impact of Paganin filter strength and unsharp masking on i) objective minimum and ii) strain mean absolute error calculated from a virtual compression study; c) mean and standard deviation of i) 1^st^ principal and ii) 3^rd^ principal strain with increasing Paganin phase retrieval strength (δ/β), with and without the addition of an unsharp mask.

To assess the effects of phase retrieval strength (δ/β) and unsharp masking on DVC performance, a 10 % virtual compression study on zero-strain images was performed. Increasing Paganin filter strength resulted in a reduction in correlation residual (objective minimum) across all tissues (Fig. 2bi), with residuals decreasing by more than an order of magnitude in AF fibres and CEP cartilage. While this would conventionally suggest improved DVC performance, displacement and strain accuracy did not follow this trend. Displacement and strain errors were minimised at low phase retrieval (typically δ/β = 2), while stronger filtering progressively increased displacement and strain errors despite lower correlation residuals (Fig. 2bii, Supplementary Fig. 2, 3). This was most pronounced for measurements of strain in the AF fibres, with mean absolute error (MAER) increased from 0.33 % (δ/β = 2) to 0.79 % (δ/β = 100).

Applying an unsharp mask substantially improved displacement and strain measurement precision and accuracy for Paganin filtered images, reducing errors while preserving the contrast enhancement from phase retrieval (Fig. 2, Supplementary Fig. 2, 3). Again, this was most pronounced for fibres, with the application of an unsharp mask reducing strain MAER from 0.79 % to 0.42 % at δ/β = 100.

#### Filtering alters interpretation of experimental strain fields

We next examined whether image filtering altered strain fields measured in response to experimental *in situ* tissue deformation. The effect was strongly tissue dependent (Fig. 2c). VEP showed little sensitivity to filtering, with mean compressive strain remaining largely unchanged. In contrast, AF fibres and CEP cartilage exhibited progressively larger principal strain magnitudes with increasing Paganin filtering. In AF fibres, mean first principal strain increased from 1.3 % in unfiltered images to 3.8 % at δ/β = 100, accompanied by greater strain heterogeneity. Unsharp masking largely eliminated these trends, indicating that strain inflation at high δ/β primarily arises from image blurring rather than contrast changes alone.

Pairwise regression of displacement and strain fields across filtering conditions further revealed tissue-dependent sensitivity (Fig. 3a, Supplementary Fig. 4, 5, 6, Supplementary Note 1). CEP cartilage showed high pointwise agreement of 3^rd^ principal strain fields between filtering conditions (median R²=0.95, Fig. 3aii), consistent with its relatively large structural feature size (autocorrelation length of 15.6 voxels). VEP pointwise linear regression of 3^rd^ principal strain had a median R² of 0.78, increasing to 0.94 after spatial averaging over 50 voxel bins. AF fibres showed substantially lower pointwise agreement of 3^rd^ principal strain (median R² = 0.66, Fig. 3ai), reflecting greater sensitivity of fine-scale strain measurements to filtering, consistent with their smaller feature size (autocorrelation length 3.5 voxels).

**Figure 3.**
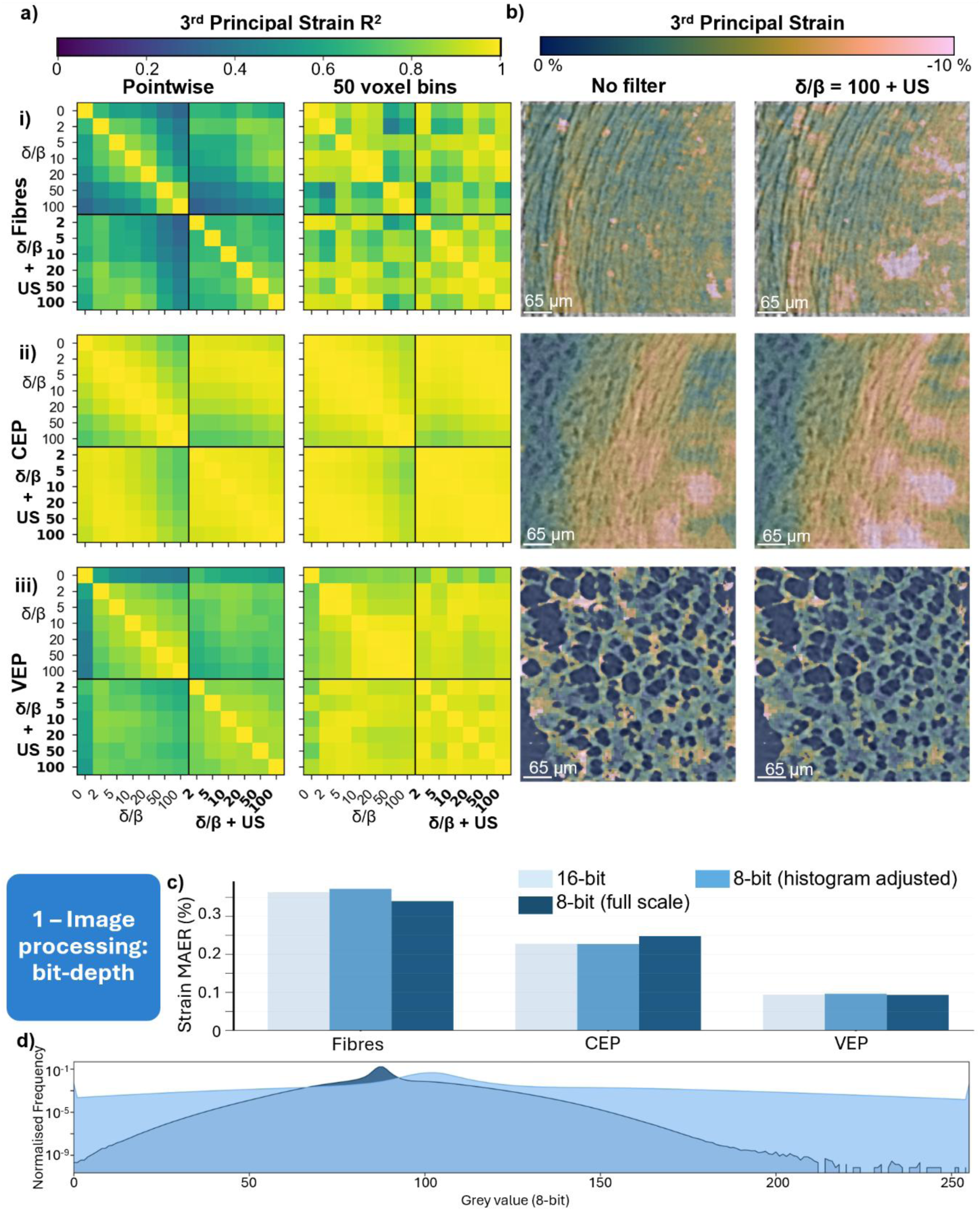
The impact of phase retrieval and image filtering on experimental strain measurements. a) R^2^ heatmaps between 3^rd^ principal strain fields measured using different filter values calculated pointwise and spatially averaged over 50 voxel bins, b) 3^rd^ principal strain maps generated from images with no filter and images with a Paganin δ/β = 100 and unsharp mask (US) for i) annulus fibrosus fibres, ii) hyaline cartilage in the CEP, iii) mineralised tissues in the VEP. The impact of bit-depth on DVC accuracy: c) Strain MAER for fibres, CEP, and VEP when performing DVC on 16-bit, histogram adjusted 8-bit, and full-range 8-bit images, d) 8-bit histograms for the histogram adjusted and full scale conversion images.

Displacement-field agreement was generally higher than strain-field agreement, although sensitivity varied between displacement components and tissues; the v component showed comparatively low pointwise agreement in both fibrous and cartilaginous regions, likely due to directional imaging artefacts (Supplementary Fig. 4, Supplementary Note 1). Filter-induced variability substantially affects pointwise strain estimation, particularly in fibrous tissue, which had a median R² value of 0.37 for 1^st^ principal strain. Spatial averaging mitigates these differences, increasing displacement and strain-field agreement (Fig. 3a), with VEP showing the greatest robustness, with 1^st^ principal strain median R² increasing from 0.76 for pointwise comparisons to 0.89 after averaging over 50 voxel bins. Fibrous tissue showed the greatest sensitivity, with 1^st^ principal strain median R² increasing from 0.37 for pointwise comparisons to 0.70 after averaging over 50 voxel bins (Supplementary Note 1).

Across all tissues, comparisons within the same processing family (Paganin-only or Paganin + unsharp mask) showed higher agreement than comparisons between families, indicating that unsharp masking changes the spatial distribution of strain rather than simply shifting strain magnitude. The improved agreement after regional averaging further suggests that image filtering primarily affects strain measurements near the spatial resolution limit of DVC.

Together, these results show that image-processing choices can directly alter both the magnitude and spatial distribution of measured strains, and therefore the biological interpretation of deformation experiments. DVC optimisation should therefore be considered not only an accuracy problem, but also a reproducibility problem.

Image filtering can be extremely useful for reducing noise in images and improving the results of processing steps such as segmentation. For example, in our previous study, higher levels of phase retrieval (Paganin δ/β = 100) were required to enable thresholding of mineralised from non-mineralised tissues in the vertebral endplates [8]; unsharp masking was used to minimise the negative effects of this on DVC accuracy. Paganin phase retrieval in multi-material systems with differing attenuation coefficients can create additional image artefacts as the assumption of a homogeneous material breaks down. This creates further difficulties for performing DVC at material interfaces. Advanced iterative phase retrieval approaches, such as Eikonal phase retrieval [23], can help reduce these artefacts and will likely offer significant benefits for performing DVC at material interfaces.

Median filtering has been shown to increase strain errors in some systems, whereas non-local-means denoising appeared to reduce strain error [15]. In general, and as demonstrated by the results of this study, filters which blur the image will result in reduced DVC accuracy due to a reduction in trackable image features. This means that simple noise reduction filters such as median, Gaussian, and mean should be avoided. Filters designed to maintain image features, such as non-local-means denoising [24] and wavelet denoising [25] are therefore better suited for DVC pre-processing.

Similarly, artificial intelligence (AI) based denoising methods should be used with caution; the same filtering process must be used between reference and deformed images, as features removed in one image but not the other will lead to high strain errors. Filtering choices are particularly important for images with small feature sizes.

#### Effect of image bit depth and intensity mapping on DVC measurements

Imaging data are often down-sampled from 16-bit to 8-bit to reduce storage and computational cost, but the effect on DVC accuracy is unclear. We compared DVC performance using 16-bit reconstructions and two 8-bit conversion strategies: histogram-adjusted conversion, in which image intensities were rescaled to utilise the full 8-bit dynamic range prior to quantisation, and full-range conversion, in which the entire 16-bit intensity range (0–65535) was directly mapped to 8-bit values (0–255) (Fig. 3d).

Virtual compression analysis showed that histogram-adjusted 8-bit images produced strain errors that were very similar to the corresponding 16-bit datasets. Across the three tissues, tensor strain MAER changed by only −0.2% to +2.4%, while SDER changed by −0.4% to +3.5% (Fig. 3c). In contrast, full-range conversion produced greater and tissue-dependent changes. The largest reduction in accuracy occurred in the lower-contrast CEP cartilage, where tensor MAER increased by 8.9% and SDER by 16.5% relative to the 16-bit analysis. Component-wise analysis showed that this was predominantly associated with the z-directional strain component, for which MAER increased by 17.5% (+445 με) and SDER by 61.7% (+1192 με) (Supplementary Fig. 7), representing a loss of ability to track the applied deformation. In contrast, full-range conversion slightly reduced tensor MAER and SDER in fibres (−6.5% and −7.7%, respectively) and had negligible effect in VEP (−0.6% and +0.5%, respectively), demonstrating that the effect of intensity mapping is strongly tissue dependent.

Regression analysis of experimental deformation fields showed excellent agreement between 16-bit and 8-bit displacement fields (R² > 0.92, Supplementary Figure 8), while strain fields remained highly consistent following histogram-adjusted conversion (R² = 0.78–0.98, Supplementary Figure 9).

These results demonstrate that DVC accuracy is largely insensitive to nominal bit depth when image intensities are appropriately rescaled before quantisation. Image contrast utilisation, rather than bit depth itself, is therefore the dominant factor governing measurement fidelity. Extra care must be taken when down-sampling images with low contrast-to-noise ratios. Additionally, appropriate rescaling may not be possible for images with separate distinct intensity peaks, for example the bimodal distribution seen when performing microCT on metal implants in bone [26], so extra caution should be taken when down-sampling.

### 2 Point cloud creation

#### Microstructurally tailored points clouds enable multiscale strain quantification

The method used to generate the DVC point cloud should be driven by the microstructure of the material of interest (Fig. 4a,b). An excellent example of this is fibrous materials. The mechanics of fibre-based materials is of interest at both the fibre [2, 4] and bulk material scales [2, 14, 27, 28], and the use of a fibre-based point cloud (Fig. 4 biii) can enable multiscale strain calculation [2]. However, the utilisation of a non-uniform point cloud may impact the measured strain field.

**Figure 4.**
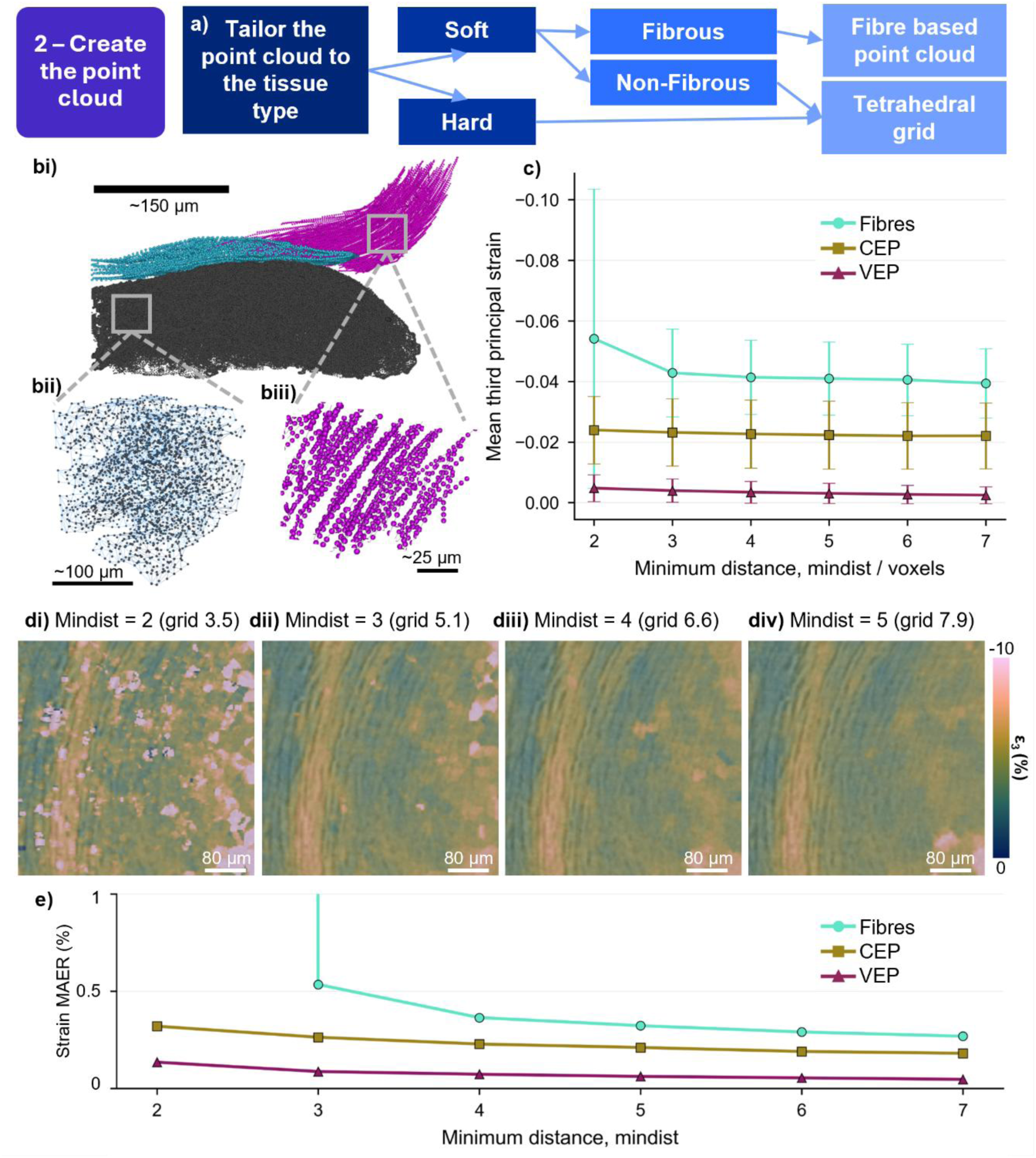
Point cloud creation; a) flow-chart for choosing point cloud generation method, b) point cloud generation across the AF-CEP-VEP junction, bii) tetrahedral grid based point cloud in the VEP, biii) fibre based point cloud in the AF; c) mean and standard deviation of experimentally measured 3^rd^ principal strain with increasing distance between points in the point cloud; d) 3^rd^ principal strain fields in the AF measured with point clouds generated using i) a minimum distance between points of 2 voxels (equivalent to a regular grid spacing of 3.5), ii) a minimum distance of 3 (equivalent to a regular grid spacing of 5.1), iii) a minimum distance of 4 (equivalent to a regular grid spacing of 6.6), iv) a minimum distance of 5 (equivalent to a regular grid spacing of 7.G); e) mean absolute strain error across fibres, CEP, and VEP with increasing distance between points.

To investigate the impact of using a fibre-based point cloud versus a regular grid for tissue-level strain measurements, density matched point clouds were generated using the fibre-based method and a regular cubic grid (Supplementary Figure 10). Voxel-wise and spatially averaged regression analysis was performed between the results interpolated and resampled to the same voxel size as the images (Supplementary Tables 2,3). Voxel-wise comparison between the fibre-derived and regular-grid strain fields yielded R² values of 0.684 (1^st^ principal strain) and 0.776 (3^rd^ principal strain).

Following spatial averaging, agreement increased substantially, reaching R² = 0.976 and 0.995 at 10-voxel cubes, indicating that the fibre-based representation preserves the strain field at biomechanically relevant length scales, enabling confident derivation of both tissue-and fibre-level strain from the same point cloud.

Using a tetrahedral grid with specified nodal spacing of segmented microstructure ensures points are placed approximately uniformly throughout complex structures without missing out thin (<10 voxel diameter) structures e.g. cartilage septae [7, 8] (Fig. 4bii). If the desired DVC point spacing is significantly lower than the minimum width of structures, then a regular grid could also be used for point cloud generation. The finite element mesh must be produced in a way that results in approximately uniform point spacing and ensures a minimum distance between points, as points placed too close together will lead to high strain errors.

#### Effect of point cloud density on the measured strain field

Increasing the density of DVC measurement points enables an increase in strain field sampling density and improved spatial resolution, without impacting displacement accuracy or DVC residuals (Supplementary Fig. 11,12). Increasing the distance between points results in a gradual reduction of measured strain magnitude (Fig. 4c), progressive strain field smoothing (Fig. 4d), and a reduction in strain error (Fig. 4e, Supplementary Fig. 11,12). The impact of point cloud density varied by tissue type; for AF fibres, a minimum distance below 3 voxels resulted in unacceptably high strain error with clear outliers in the experimentally measured strain field (Fig. 4di). Parameter sets were considered acceptable when the absolute strain error was below 10% of the characteristic experimental strain magnitude; for the 10% virtual-compression test, this corresponded approximately to an absolute strain-error threshold of 1 % strain. For CEP and VEP tissues, acceptable strain accuracy was achieved even when the highest tested density (minimum distance 2 voxels) was used.

Point density should be selected according to the size of the mechanical features of interest. As an initial design guideline, point spacing should be sufficiently small to sample mechanically relevant structures at several locations across their width; approximately five points across a feature provides a useful starting point. For example, resolving strain variations within a trabecula that is 10 voxels in diameter would require a point spacing of approximately 2 voxels or less.

### 3 Running DVC analysis

#### Time costs

Subvolume size, number of points in the point cloud, and correlation procedure all come with associated time costs (Figure 5a). In general, increasing the accuracy and spatial resolution of DVC analysis will lead to an associated increase in computational time. For small point clouds consisting of up to a few thousand points, the time cost for DVC is negligible and there is no need to compromise accuracy or spatial resolution for the sake of time. For point clouds containing several million points, computational time can become a major bottleneck, and compromises may need to be made.

**Figure 5.**
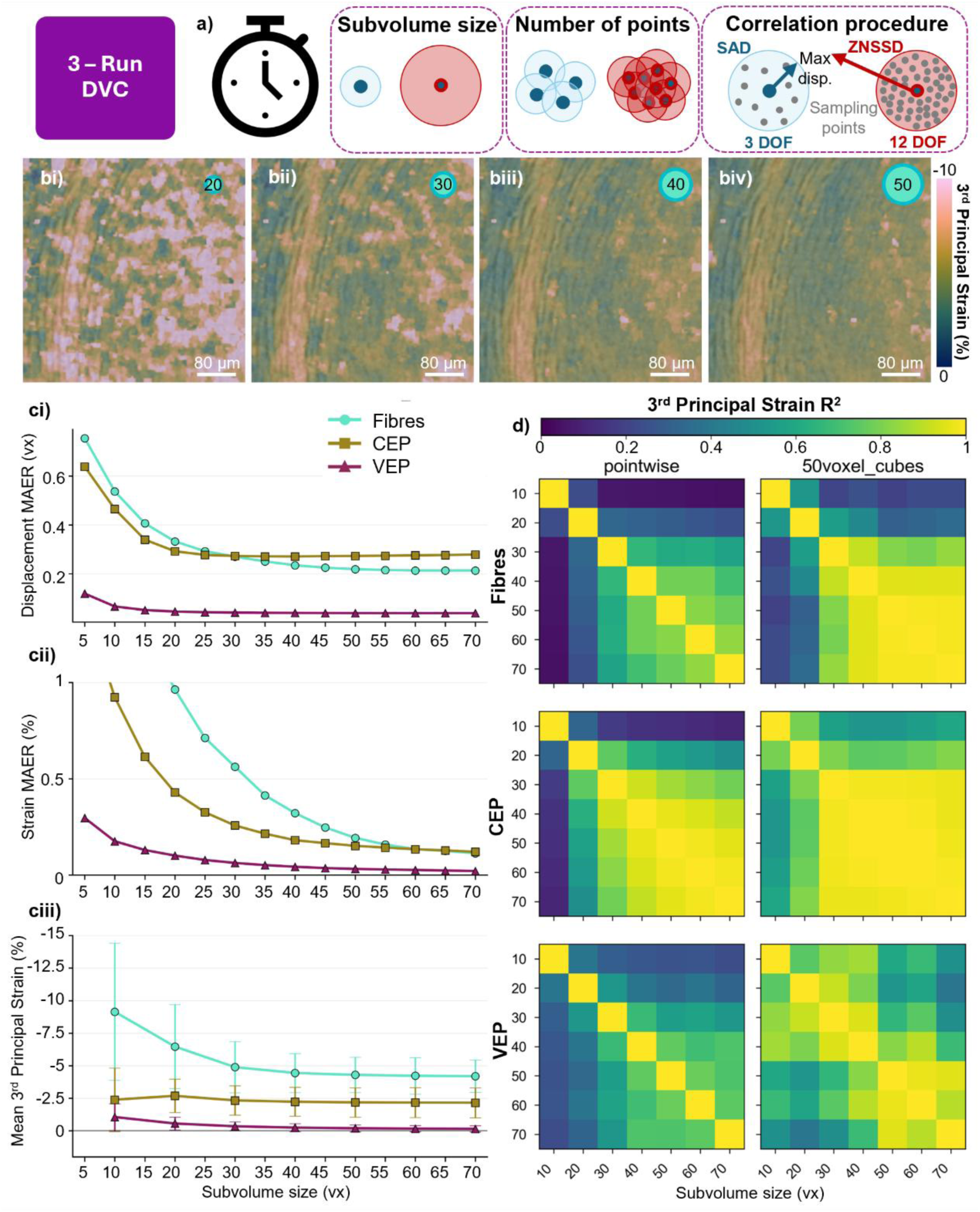
Running DVC. a) Summary illustration of parameters affecting computational time required for analysis; b) 3rd principal strain fields measured using different subvolume diameters of i) 20 voxels, ii) 30 voxels, iii) 40 voxels, iv) 50 voxels; c) the impact of increasing subvolume size on i) displacement MAER, ii) strain MAER, and iii) 3^rd^ principal strain; d) pointwise and 50 voxel binned linear regression analysis of 3^rd^ principal strain values measured using different subvolume sizes in AF fibres, CEP cartilage, and VEP mineralised tissues.

Performing initial low-resolution DVC analysis across large areas to identify regions for subsequent high-resolution analysis is one potential strategy for reducing the computational burden of analysis. However, it must be noted that the measured strain magnitudes cannot be compared between analyses performed at different spatial resolutions. As shown in the previous sections, point cloud density can have a significant effect on measured strain magnitude.

Reducing the number of sampling points placed within each subvolume can increase computational speed without affecting spatial resolution. Displacement and strain accuracy generally increase with increasing sampling points, and low-contrast materials (Fibres and CEP) are more affected than those that comparatively have higher contrast (VEP) (Supplementary Figures 14, 15).

#### The effect of subvolume size on DVC

Increasing subvolume size had a strong and tissue-dependent effect on DVC accuracy in the virtual compression study. Across all tissues, displacement and strain errors decreased with increasing subvolume size (Fig. 5c, Supplementary Fig. 16, 17), reflecting improved correlation robustness and reduced noise when larger image regions were used for tracking.

The magnitude of this effect differed between tissue types (Fig. 5c). AF fibres showed the highest sensitivity to subvolume size, with strain MAER from a 10 % virtual compression test greater than 1 % for subvolume diameters below 20 voxels, with a plateau in strain error appearing at around 60 voxels. In contrast, mineralised VEP tissues maintained strain MAER more than an order of magnitude below the applied strain even at a subvolume diameter of 5 voxels, and minimal improvement in strain accuracy beyond 30 voxels. CEP showed an intermediate response.

The real experimental strain-field analysis showed the expected smoothing effect of increasing subvolume diameter (Fig. 5b). In general, measured strain magnitude decreased with increasing subvolume size. This demonstrates that larger subvolumes suppress local strain heterogeneity and reduce apparent strain magnitude, consistent with spatial averaging of the measured deformation field. Linear regression further showed that displacement fields were highly consistent across subvolume sizes (Supplementary Fig. 18), whereas strain-field agreement was weaker at the pointwise level but improved after spatial averaging (Fig. 5d, Supplementary Fig. 19, 20), indicating that subvolume-dependent differences primarily arose from local strain fluctuations. Together, these results demonstrate that smaller subvolumes preserve fine strain gradients but are more susceptible to correlation noise, whereas larger subvolumes improve precision at the expense of attenuating biologically relevant strain heterogeneity.

### 4 Calculating the strain field

#### Impact of strain window size on the strain field

In iDVC, the strain tensor is estimated at each point from a cloud of neighbouring points using a least-squares fit to a second-order Taylor series expansion of the displacement vector field [29, 30]. In this, the number of points used for fitting can be varied to obtain different levels of strain field smoothing.

Increasing strain window size from 10 to 75 points resulted in substantial strain field smoothing (Fig. 6b). The minimum required strain window size for suitable strain accuracy (MAER < 1 %) varied by tissue type, with VEP requiring 20 points, CEP 25 points, and AF fibres 30 points (Supplementary Fig. 21).

**Figure 6.**
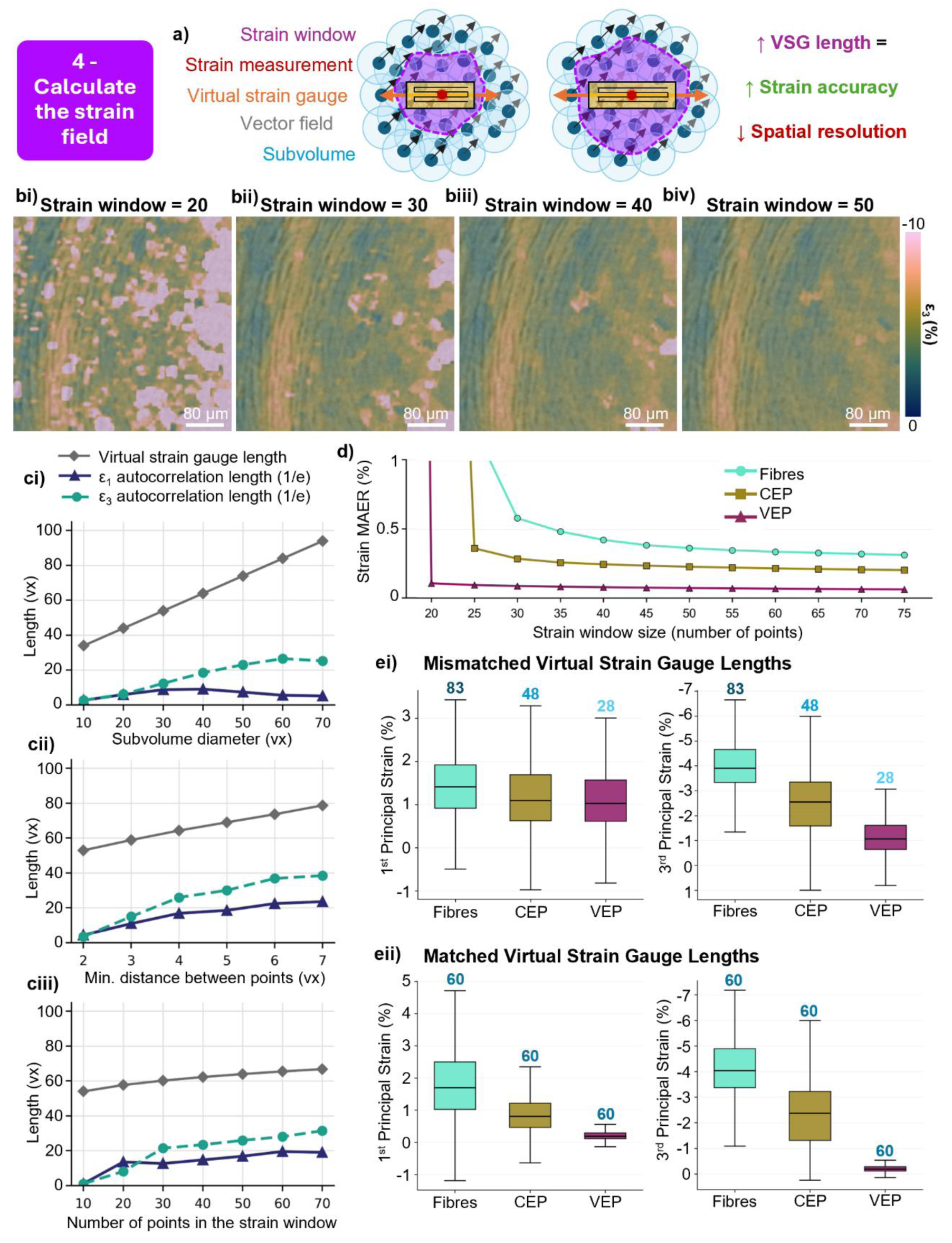
Calculating the strain field. a) illustration of the virtual strain gauge; b) 3^rd^ principal strain fields calculated using different strain windows of i) 20 points, ii) 30 points, iii) 40 points, and iv) 50 points; c) 3D autocorrelation length (1/e) and virtual strain gauge length calculated from the experimental 1^st^ and 3^rd^ principal strain fields in the annulus fibrosus fibres with increasing i) subvolume size, ii) point spacing, and iii) strain window size. d) strain MAER with increasing strain window size; e) The effect of mismatched virtual strain gauge lengths on measured load sharing between materials, demonstrated by boxplots of 1^st^ and 3^rd^ principal strain in fibres, CEP, and VEP for i) mismatched virtual strain gauge lengths and ii) matched virtual strain gauge lengths; numerical labels indicate the virtual strain gauge length in voxels.

Smoothing of the displacement field is vital to obtaining high-resolution strain fields. Strain is calculated from the gradient of the displacement field, and so any noise in the displacement field will be amplified during the strain calculation if not smoothed out. Fitting a Taylor series expansion [29] or polynomial [2] to the displacement field is a robust and flexible way to do this. Other methods include filtering the displacement field prior to strain calculation [28], which removes high displacement gradients, or employing regularisation techniques [31], which use material property assumptions to prevent the measurement of erroneously high strains due to image noise.

#### Optimising the spatial resolution of DVC

The spatial resolution of the measured strain field depends on three parameters: the size of subvolumes used for correlation, the distance between measurement points (subvolume overlap), and the level of smoothing used during the strain calculation.

Depending on the scientific questions of interest, it is often useful to obtain the highest possible resolution strain fields from an in situ imaging dataset. The highest achievable resolution depends strongly on the image properties of the material of interest.

One method to quantify the spatial resolution of DVC is to calculate the virtual strain gauge length (Fig. 6a) – the span of points over which strain is being calculated. This can be approximately defined as:

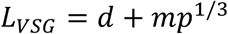

Where *d* is the subvolume diameter, *m* is the distance between measurement points (subvolume centres), and *p* is the number of points used for fitting during strain calculation. This is an approximate effective measurement length, and is not exact for irregularly distributed points.

Autocorrelation analysis of 1^st^ and 3^rd^ principal strain fields in the AF demonstrated that all three parameters altered the spatial resolution of the resulting strain fields (Fig. 6c). Increasing subvolume diameter has the greatest impact on virtual strain gauge length, followed by point spacing, then strain window size. Autocorrelation analysis showed that the characteristic spatial variation of the measured strain field can occur over length scales smaller than the nominal virtual strain gauge length, indicating that the gauge length should not be interpreted as a sharp lower bound on detectable strain-field structure.

These results provide a useful guide for the order in which to optimise DVC parameters to achieve the highest spatial resolution of the strain field:

1. Identify the smallest possible subvolume size that gives accurate strain measurements (in general, strain error should be below 10 % the experimentally measured strain), while keeping point spacing and strain window size large.
2. Find the minimum distance between points which gives acceptable strain accuracy whilst using the identified minimum subvolume size and keeping the strain window size large.
3. Reduce the number of points in the strain window until the strain accuracy threshold is reached.

#### Strain fields must be compared at matched spatial resolution across materials

Our analysis shows that any parameter contributing to the virtual strain gauge (VSG) length influences both the apparent magnitude and spatial heterogeneity of experimentally measured strain fields. As DVC accuracy is strongly governed by image texture and contrast, different materials impose different limits on the spatial resolution at which strain can be reliably measured. Across all analyses in this study, the highly mineralised vertebral endplate (VEP) consistently supported accurate strain measurements at smaller VSG lengths than the AF and CEP, owing to its higher image contrast and strong correlation texture.

Although maximising spatial resolution is generally desirable in DVC, strain fields should be compared across materials only when measured at equivalent VSG lengths. When VSG lengths were not matched between tissues (Fig. 6e,i), the relative strain partitioning across the intervertebral disc differed markedly from that obtained using matched spatial resolution (Fig. 6e,ii). For compressive strain (third principal strain), ratios were as follows:

- VSG mismatched conditions: Fibres: CEP: VEP = 3.1 : 1.9 : 1
- VSG matched conditions: Fibres: CEP: VEP = 14 : 7.7 : 1 Similarly, for tensile strain (first principal strain):
- VSG mismatched conditions: Fibres: CEP: VEP = 1.2 : 1 : 1
- VSG matched conditions: Fibres: CEP: VEP = 6.3 : 3 : 1

These large shifts arise because increasing VSG length spatially averages local strain gradients, suppressing peak strains and reducing apparent heterogeneity.

Consequently, mismatched spatial resolution can substantially distort the inferred load-sharing behaviour between neighbouring materials.

These findings highlight an important consideration for DVC studies of heterogeneous or multi-material systems: differences in measured strain may reflect differences in spatial resolution as much as intrinsic mechanical behaviour. Matching VSG length across materials is therefore essential for robust quantitative comparison and for avoiding erroneous interpretation of strain transfer across material interfaces.

### 5 Strain tensor analysis

Optimising DVC for heterogeneous materials requires consideration not only of displacement tracking accuracy, but also of how displacement fields are converted into physically meaningful strain metrics. The AF has a complex composite structure, with circumferential collagen-rich lamellae at alternating orientations encircling the nucleus pulposus [2]. Considered directional analysis of strain tensors can provide greater insight into load transfer within this complex system compared to analysis of strain magnitude alone.

Principal strains provide an intuitive, coordinate-independent description of the dominant tensile and compressive deformation modes, and visualisation of the first and third principal strains in annulus fibrosus (AF) tissue revealed highly heterogeneous local strain patterns arising from the fibrous lamellar architecture [2, 27] (Fig. 7a,b).

**Figure 7.**
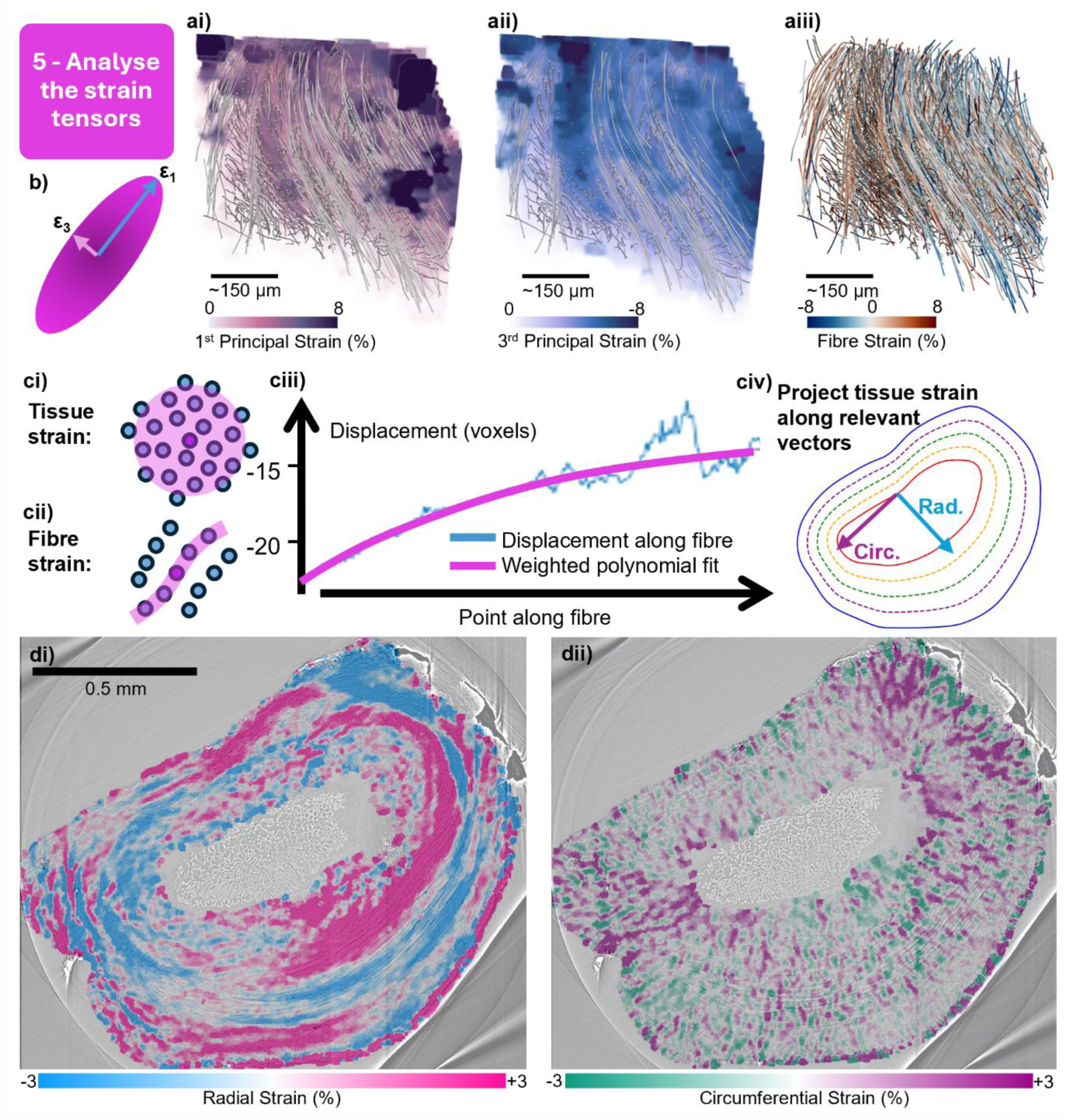
Analysing the strain tensors. a) 3D renderings of i) 1^st^ principal, ii) 3^rd^ principal, and iii) fibre strain in the AF fibres; b) illustration of 1^st^ and 3^rd^ principal strain vectors; c) illustrations of i) tissue-level strain calculation and ii) fibre strain calculation, iii) example of how a weighted polynomial fit can be used to smooth out noise in the displacement field, iv) example of anatomically relevant vectors (radial and circumferential) in the IVD; di) radial and ii) circumferential strain fields in the annulus fibrosus of the IVD reveal microstructurally driven strain patterns.

However, while principal strains effectively describe the magnitude of local deformation, they do not explicitly indicate whether deformation occurs along structurally meaningful directions such as collagen fibres or anatomical axes. This limits their interpretability in highly anisotropic materials, where mechanical behaviour is strongly governed by underlying microstructure. Quantifying the direction of principal strain vectors with respect to fibre orientation [2] or the experimental loading axis [8], can provide further information on deformation mechanisms.

Strain calculation must also be considered in relation to spatial scale and material architecture. Conventional tissue-level strain estimation uses displacement gradients calculated from neighbouring DVC points, with the effective smoothing determined by the strain window (Fig. 7ci). For fibrous materials, multiscale strain analysis can provide a more appropriate framework by enabling strain to be calculated directly along fibres (Fig. 7cii). In our previous studies of intervertebral disc mechanics [2], fibre strain was quantified by applying polynomial fits to displacement along the fibre direction, enabling smoothing of noisy displacement fields (Fig. 7ciii). This enabled strain to be evaluated across multiple structural length scales, revealing deformation modes that were obscured using tissue-level strain tensors alone. Projecting strain tensors along anatomically relevant directions can further improve mechanical interpretation. In the intervertebral disc, radial and circumferential strain maps revealed clear microstructure-driven deformation patterns that were not readily apparent from principal strain maps alone [2] (Fig. 7civ,d).

These findings demonstrate that strain analysis should be adapted to both the material architecture and the physical scale of deformation. As a practical guide, we recommend that principal strains are used as default descriptors for isotropic or weakly anisotropic materials; multiscale or fibre-aligned strain metrics are used for fibrous materials; and tensor projection onto anatomically relevant axes is used when organ-scale structural loading is of interest. Such structure-informed strain analysis enables DVC to move beyond generic strain mapping towards mechanistically meaningful interpretation in complex multi-material systems.

## Conclusions

Digital volume correlation provides a powerful means of investigating deformation in heterogeneous materials, but the measured strain field is not independent of the analytical choices used to generate it. Across fibrous, cartilaginous and mineralised tissues, we show that image processing, point-cloud architecture, subvolume size and strain calculation each influence the accuracy, spatial resolution and apparent magnitude of local strain. These effects depend strongly on material microstructure and image characteristics: fine-scale fibrous structures are particularly sensitive to image blurring from filtering, whereas low-contrast materials are more vulnerable to inappropriate intensity rescaling during bit-depth reduction. Importantly, improved correlation residuals do not necessarily indicate improved displacement or strain accuracy, demonstrating that optimisation should be based on validation against known deformation rather than residual minimisation alone.

The spatial scale of strain measurement emerges as a central determinant of both measurement accuracy and mechanical interpretation. Increasing subvolume size, point spacing or strain-window size improves robustness but progressively smooths local strain heterogeneity, with subvolume size exerting the strongest influence on the effective measurement scale. We therefore recommend a sequential optimisation strategy in which the smallest reliable subvolume is identified first, followed by optimisation of point spacing and strain-window size against a predefined strain-error threshold – generally 10 % of the measured strain. Microstructurally informed point clouds and anatomically relevant strain metrics can further improve interpretation by enabling deformation to be evaluated at the structural scales and along the directions most relevant to the material.

Crucially, meaningful comparisons between materials require strain fields to be evaluated at matched spatial resolution. In the intervertebral disc, adjusting the virtual strain gauge length changed the apparent relative partitioning of compressive strain between fibrous, cartilaginous and mineralised tissues from approximately 3:2:1 to 14:8:1, illustrating how mismatched measurement scales can fundamentally alter interpretations of load transfer.

An overview of the key steps and considerations for DVC study design, optimisation, and analysis is given in Figure 8. Although the precise parameter values identified here are specific to the imaging conditions and materials examined, the underlying framework is broadly applicable to biological tissues, engineered composites and other heterogeneous systems. By integrating image-characteristic assessment, deformation-based validation and explicit control of measurement scale, this approach provides a practical basis for more accurate, reproducible and mechanically meaningful DVC studies.

**Figure 8.**
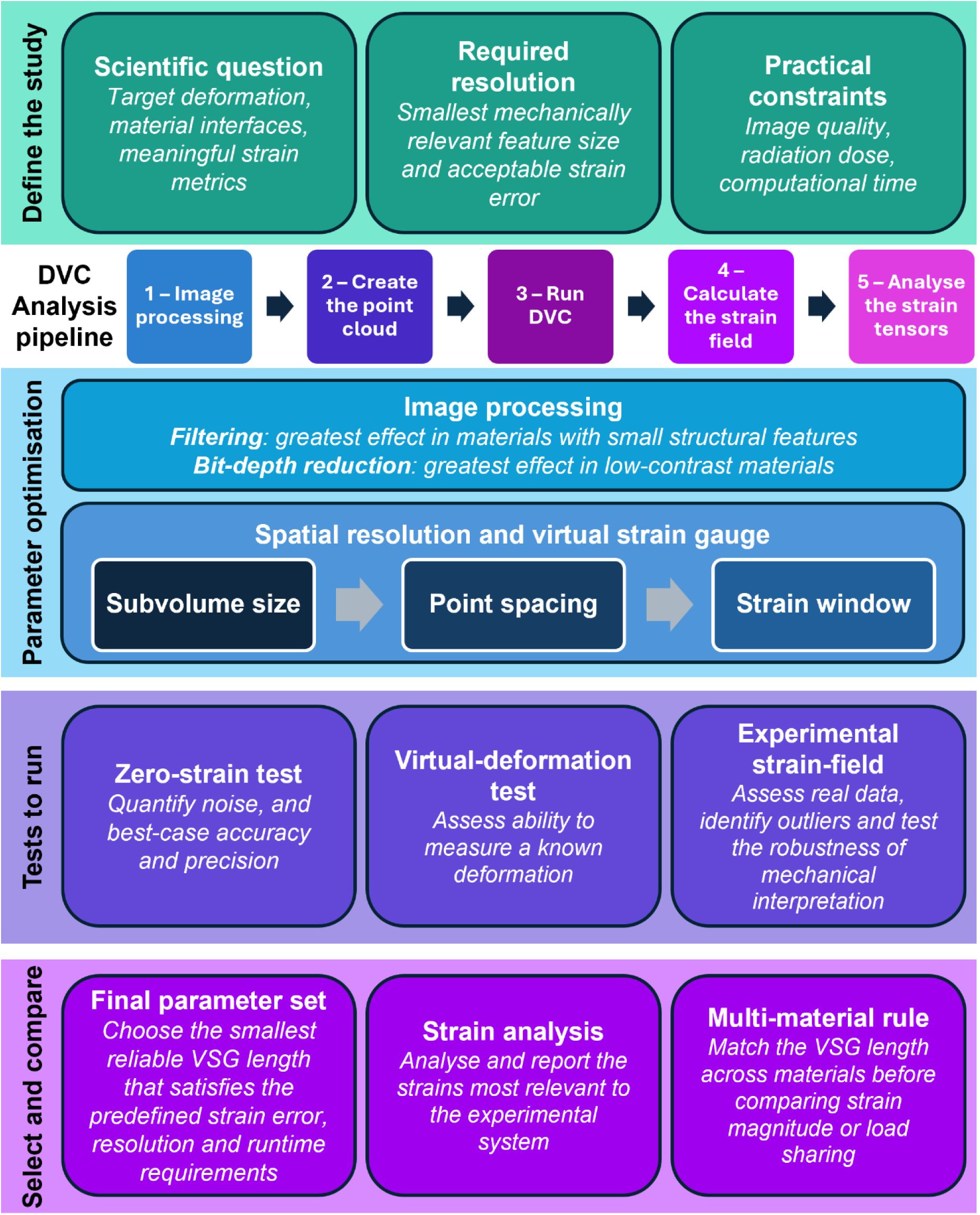
Overview schematic of DVC study design, optimisation, and analysis. The scientific question, required spatial resolution and practical constraints should first be defined. The workflow then proceeds through image processing, point-cloud generation, DVC, strain-field calculation and strain-tensor analysis. Image-processing parameters should reflect material-specific image characteristics: filtering has the greatest effect in materials with small structural features, whereas bit-depth reduction particularly affects low-contrast materials. Subvolume size, point spacing and strain-window size collectively determine the accuracy and effective spatial resolution of the displacement and strain measurements. Candidate parameter sets should be evaluated using complementary zero-strain, virtual-deformation and experimental tests. Zero-strain tests quantify measurement noise, and best-case precision and accuracy; virtual-deformation tests assess the recovery of a known imposed deformation; and experimental strain fields reveal outliers and test the robustness of the resulting mechanical interpretation. The final parameter set should provide the smallest reliable virtual strain gauge (VSG) satisfying predefined requirements for strain error, spatial resolution and computational time. Strain measures relevant to the experimental system should then be analysed and reported. When comparing different materials, the VSG length should be matched to avoid confounding differences in strain magnitude with differences in measurement resolution.

## Methods

### Animals and sample preparation

Male 8-week-old Sprague Dawley rats kept in controlled conditions (12:12 h light:dark cycles, 22°C and *ad libitum* access to food and water) at the University of Manchester Biomedical Sciences Facility with all tissue collection procedures performed in accordance with the UK Animals (Scientific Procedures) Act 1986, local regulations set by the UK Home Office with institutional approval obtained from the University of Manchester Animal Welfare and Ethics Review Board under the Home Office Licence (#70/8858 or I045CA465). Whole spines were dissected and snap frozen in liquid nitrogen prior to storage at −80°C. Within 24 h of *in situ* mechanical testing, samples were thawed at room temperature before further dissection.

IVD with half vertebrae either side and posterior elements removed were used for analysis. Dissections were performed using a high precision diamond cutting blade (Accumtom-50; Struers, UK). Sample damage was minimized during dissection through the use of water cooling, a slow feed speed (0.05 mm/s) and 1,800 rpm.

For virtual compression tests images from a lumbar 3-4 IVD. Experimental strain analysis was performed on the lumbar 2-3 IVD from the same rat. This data was taken from a larger study presented elsewhere [8].

### *In situ* mechanical testing

*In situ* imaging of rat intervertebral discs was performed as described previously [8]. Example imaging datasets are available here [32] (https://doi.org/10.5522/04/26789212) and here [33] (https://doi.org/10.5522/04/30286255).

Samples were mounted in a Deben CT500 (Deben, UK) rig containing a 100 N loadcell (accuracy ±1% of full scale) and custom-built open frame design [8, 9]. Custom 3D-printed (Acrylonitrile Butadiene Styrene, ABS) sample holders were used for sample mounting.

Spinal units were fixed to the bottom 3D-printed compression plate using superglue (Vetbond, 3M) and left free at the top. Samples were placed inside polyimide tubing (diameter 6 mm, wall thickness 75 μm) filled with phosphate-buffered saline (PBS) to maintain sample hydration during scanning. Samples were held under cumulative axial compression, applied in the z direction from beneath the bottom sample holder during imaging. A 1 N compressive preload was first applied, and images taken 20 min after load application to allow for stress relaxation.

Experimental strain analysis was performed on images taken at preload and after a 40 μm compressive displacement. For virtual compression tests, two images were taken after the full loading protocol was applied (1 N preload and four 40 μm displacement steps) without applying any deformation to the sample between images.

### Synchrotron X-ray computed tomography

sCT of whole IVDs was performed at the Diamond-Manchester Imaging Branchline I13-2 [34] at Diamond Light Source, UK, as described previously [8]. Imaging used a pink beam with mean energy 27.6 keV, 0.5 m sample-detector distance, and filters to remove low energy X-rays (Pyr. Graphite 0.28 mm + 1.06 mm, Al 3.2 mm, Fe 0.14 mm). Images had a 1.625 µm voxel size and 4.2 mm x 3.5 mm field of view. Limited angle tomography artefacts resulting from the open frame mechanical testing rig were avoided through the use of carbon fibre pillars which were designed to be X-ray transparent, allowing collection of usable tomographic projections at all angles.

Tomographic datasets were reconstructed using the open-source pipeline Savu [35]. Firstly, projections were pre-processed using flat and dark field correction and lens distortion correction [36], followed by ring artefact removal [37]. Datasets were then reconstructed using the filtered back projection algorithm [38] with or without Paganin phase retrieval [21] and formatted in 16-bit tif stacks.

### Image-characteristic analysis

Image characteristics were calculated from one image reconstructed with a Paganin δ/β value of 100 and an unsharp mask with radius 2.5 voxels applied to the reconstructed image [8]. Volumes of interest (VOIs) measuring 200 × 200 × 200 voxels were extracted from the annulus fibrosus (AF) and vertebral endplate (VEP), with each VOI positioned within the respective tissue to avoid tissue interfaces. To accommodate the smaller thickness of the cartilage endplate (CEP), a 200 × 200 × 70 voxel VOI was used. The CEP was manually segmented from the adjacent AF and VEP using the paintbrush tool in Avizo (2023.2, Thermo Fisher Scientific). Voxels outside the segmented CEP, including voxels belonging to the adjacent AF and VEP, were assigned an intensity value of zero before export and analysis.

Image characteristics were quantified using greyscale-intensity distributions, contrast-to-noise ratio (CNR), and three-dimensional radial greyscale autocorrelation. Intensity distributions and the mean CEP intensity were calculated using only voxels within the segmented tissue mask; zero-valued voxels outside the CEP mask were excluded from these calculations. CNR was calculated as

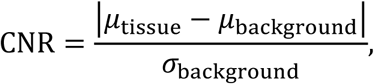

where *μ*_tissue_ and *μ*_background_ are the mean greyscale intensities within the tissue and background regions, respectively, and *σ*_background_ is the standard deviation of the background intensity. The background greyscale intensities were calculated from a 200 voxel cubed image region placed outside the sample.

Three-dimensional autocorrelation was calculated using the acf_analysis.py implementation supplied with the *Digital Volume Correlation Challenge 2.0* dataset [39, 40] (NIST repository DOI: <u>10.18434/mds2-4129</u>). For the CEP analysis, the rectangular volume supplied to the autocorrelation code retained the zero-valued voxels outside the segmented CEP.

The characteristic greyscale correlation length, l_1/*e*_, was defined as the first separation at which the radial autocorrelation decreased to 1/*e*. Additional correlation lengths defined at 0.10 and 0.01 are given in Supplementary Fig. 1.

Spatial autocorrelation was calculated for experimentally measured 1^st^ and 3^rd^ principal strain fields in the annulus fibrosus using the same methodology, to assess how parameter choices influenced the characteristic length scale of strain-field variation.

Point-based strain values were interpolated onto a regular three-dimensional grid with a voxel spacing of 1 voxel (1.625 μm) using nearest neighbours interpolation.

### DVC point cloud generation

All DVC point clouds were generated from images reconstructed using a Paganin δ/β value of 100 and an unsharp mask with radius 2.5 voxels applied to the reconstructed image [8]. Three point-cloud generation strategies were used: regular Cartesian grids, tetrahedral-mesh-derived point clouds and fibre-derived point clouds.

Regular Cartesian point clouds were generated by positioning measurement points at a uniform isotropic spacing, *d*, along the x-, y- and z-directions of the prescribed analysis volume. The grid spacing was varied as specified for each analysis.

For mineralised vertebral endplate (VEP) tissues, point clouds were generated as described previously [8]. Briefly, mineralised tissue was segmented from the reference-state image in Avizo using interactive thresholding, with the threshold selected individually for each image to distinguish mineralised tissue from the adjacent marrow space. The resulting binary mask was cleaned and smoothed before conversion into a triangulated surface. The surface was simplified using the prescribed minimum node separation and converted into a tetrahedral volume mesh. The nodes of the tetrahedral mesh were exported as the DVC point cloud.

A corresponding tetrahedral-mesh approach was used for the non-mineralised cartilage endplate (CEP). The CEP was manually segmented from the reference-state image using the Brush Tool in Avizo 2023.2 (Thermo Fisher Scientific, Waltham, MA, USA), excluding the surrounding annulus fibrosus and mineralised VEP tissues. A triangulated surface was generated from the binary segmentation and simplified using the prescribed minimum node separation. The enclosed volume was then tetrahedralised, and the surface and internal nodes of the resulting mesh were used as DVC measurement points. The target node separation was varied to investigate the effect of point-cloud density on DVC performance.

For annulus fibrosus fibres, fibre-derived point clouds were generated as described previously [2]. Briefly, individual collagen fibres were traced in the reference-state image using the XFiber module in Avizo 2023.2. Traces corresponding to reconstruction artefacts, including ring artefacts and streaks arising from mineralised tissues, were removed. The resulting fibre spatial graph was exported, and the fibre centrelines were resampled at the prescribed arc-length interval to generate x-, y- and z-coordinates along each fibre using code available here: https://github.com/DiamondLightSource/tomosaxs/tree/main/Code/CT%20%26%20DVC. These coordinates were used as DVC subvolume centres, while retaining the corresponding fibre identification number. The centreline sampling interval controls the spacing of points along an individual fibre. The minimum distance between traced fibres was varied in the Trace Correlation Lines module to impose a minimum 3D distance between points on neighbouring fibres.

To compare the global measurement densities of the regular and non-uniform point clouds, a density-equivalent isotropic Cartesian-grid spacing was calculated. Point density was defined as

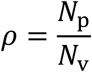

where *Np* is the number of DVC measurement points and *Nv* is the number of labelled voxels in the corresponding binary tissue mask. Binary masks were available for all tissues. All generated measurement points produced valid DVC solutions and were therefore included in *Np*. The equivalent Cartesian-grid spacing was then calculated as

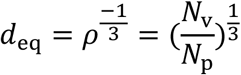

When the tissue volume is expressed as a labelled voxel count, *deq* is reported in voxels. For an isotropic voxel size *a*, the corresponding physical spacing is

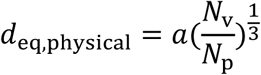

This quantity represents the spacing of a cubic Cartesian grid with the same mean number of measurement points per unit tissue volume (Supplementary Fig. 13). It does not represent the mean nearest-neighbour distance or the local spacing within the irregular tetrahedral- and fibre-derived point clouds.

### Digital Volume Correlation Analysis

All digital volume correlation analysis was performed using iDVC (24.1.0, https://tomographicimaging.github.io/iDVC). All analysis was run using spherical sub-volumes, 12 degrees of freedom, zero-normalised sum-of-squared difference objective function, and tricubic interpolation. A list of the values (subvolume size, sampling points, displacement maximum, strain window size) used for each DVC run is provided in Supplementary Table 1.

Lagrangian strains were calculated from the measured displacement fields using the strain executable distributed with iDVC. The calculation follows the three-dimensional formulation described by Bay [29] and the second-order discrete-displacement approach of Geers *et al.* [30]. At each DVC measurement point with material coordinate **X**, the *p* nearest neighbouring points defined the strain window. A second-order Taylor polynomial was fitted to the three-dimensional displacement field by ordinary least squares:

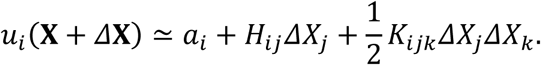

Here, *u_i_* is a component of the displacement vector, *Δ***X** is the reference-position offset of a neighbouring point from the central point, *a_i_* is the fitted local displacement, *H_ij_* = *∂^2^u_i_*/ *∂X_j_* is the material displacement gradient and *K_ijk_* = *∂*^2^*u_i_*/(*∂X_j_ ∂X_k_*) contains the second spatial derivatives; repeated indices imply summation. Including the quadratic coefficients allows the local displacement gradient to vary across the fitting window.

The window size *p* controls the balance between noise suppression and spatial resolution: larger windows provide greater smoothing but average over a larger region. For the strain-window sensitivity analysis, *p* was varied from 10 to 75; the value used for every DVC run is given in Supplementary Table 1. No DVC measurement points were excluded before fitting.

The deformation gradient **F** was obtained from the fitted displacement gradient as

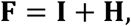

where **I** is the identity tensor. The Green–Lagrange strain tensor **E** was then calculated as

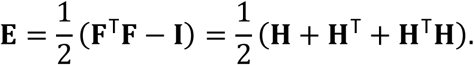

Because the derivatives are taken with respect to the undeformed reference coordinates, **E** is a finite Lagrangian strain measure. The six independent components (*E_xx_*, *E_yy_*, *E_zz_*, *E_xy_*, *E_xz_* and *E_yz_*) were retained; the off-diagonal components are tensor shear strains. Principal strains and directions were obtained by eigen decomposition of the symmetric strain tensor, with the eigenvalues ordered *E*_1_ ≥ *E*_2_ ≥ *E*_3_.

The virtual strain-gauge (VSG) length was defined as

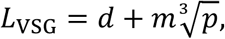

where *d* is the spherical subvolume diameter, *m* is the centre-to-centre distance between DVC sampling points and *p* is the number of points in the strain window. This measure captures the combined spatial averaging introduced by the correlation subvolume, sampling density and strain fitting.

### Virtual-deformation analysis

Virtual deformation analysis was performed using two independent images taken of the sample in the same state, i.e. zero strain between the two images. The second of these images was then manipulated in Avizo by opening the crop editor and changing the voxel size in z from 1 to 0.9. This manipulated image was then resampled back to a 1×1×1 grid using Lanczos resampling. This resulted in a uniform axial displacement gradient of *∂u_z_*/ *∂Z* = −0.10, with all other displacement-gradient components equal to zero. Two images used for virtual compression analysis (8-bit, Paganin δ/β = 2) are available in the supplementary dataset (https://doi.org/10.5522/04/33733111) [41].

Strain and displacement errors were evaluated at all *N* valid DVC measurement points. The subscript *k* denotes a measurement point, *c* denotes one of the six independent strain components (*E_xx_*, *E_yy_*, *E_zz_*, *E_xy_*, *E_xz_*, *E_yz_*), and *j* denotes one of the three displacement components *u*, *v* and *w*. Their component-index sets are denoted C*_E_* and C*_u_*, respectively.

The virtual deformation imposed a uniform engineering compression of 10% in the *z* direction, corresponding to an axial stretch of *λ_z_* = 0.90, with no lateral deformation or shear. The target Green–Lagrange strain was therefore

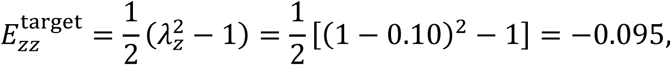

with the six target components, ordered as xx, yy, zz, xy, xz and yz, given by

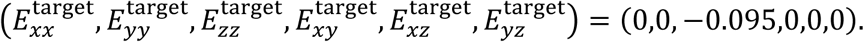

For displacement, let *z*_0,*k*_ be the original *z* coordinate of point *k*, and let *z*_ref_ be the coordinate of the fixed reference plane used to generate the virtually compressed image. Defining

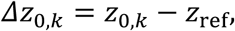

the target displacement vector at point *k* was

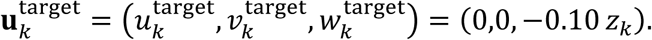

The signed strain error for component *c* at point *k* was defined as the measured-minus-target difference

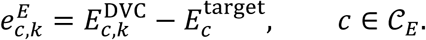

The signed displacement error was defined analogously as

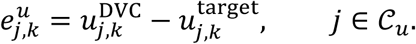

Thus, displacement error is the difference between the DVC-measured and prescribed displacement for each Cartesian component at each measurement point. The corresponding absolute errors were

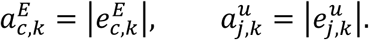

The tensor-level mean absolute strain error was calculated by pooling the absolute errors from all six independent strain components and all *N* points:

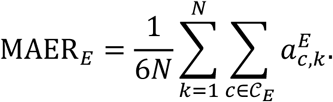

The strain SDER was the sample standard deviation of those same 6*N* absolute component errors:

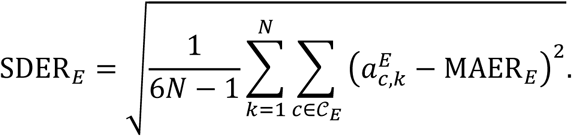

The corresponding mean absolute displacement error was calculated by pooling the absolute errors from all three displacement components:

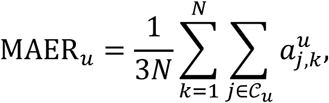

and the displacement SDER was

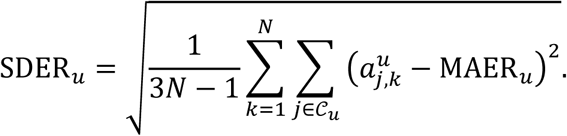

Here, MAER describes the mean magnitude of the component-wise error, whereas SDER describes the spread of the component-wise absolute errors. The 6*N* − 1 and 3*N* − 1 denominators apply Bessel’s correction and correspond to a sample standard deviation. MAER was used to describe accuracy, and SDER precision, as described by Liu and Morgan [42].

### Image processing and phase retrieval

To assess the influence of image processing on DVC performance, tomographic datasets were reconstructed using a range of Paganin single-distance phase-retrieval strengths. The ratio of the refractive index decrement to the absorption index, δ/β, was set to the values listed in Table 1, together with a reconstruction without phase retrieval. Phase retrieval was performed using Savu [35], with an assumed X-ray energy of 27.6 keV and propagation distance of 0.5 m.

**Table 1.** Paganin δ/β value and corresponding unsharp mask radius applied to the reconstructed image.

| $\delta/\beta$ | 2 | 5 | 10 | 20 | 50 | 100 |
| --- | --- | --- | --- | --- | --- | --- |
| Unsharp mask radius (voxels) | 0.5 | 0.8 | 1 | 1.3 | 1.5 | 2.5 |

For each phase-retrieval setting, additional image volumes were generated with and without unsharp masking applied to the reconstructed image. The unsharp mask was implemented using ImageJ [43] with a Gaussian radius corresponding to the values in Table 1, and a mask weight of 0.9. Identical processing parameters were applied to the reference and deformed volumes within each image pair.

DVC analyses were repeated for all filtering conditions while maintaining constant point-cloud coordinates and correlation parameters. Objective minimum, displacement error and strain error were quantified using the virtual-deformation dataset, and the effect of image processing on experimentally measured displacement and strain fields was evaluated using the corresponding experimentally loaded scan pair.

## Statistical Analyses

### Regression analysis of displacement and strain fields

Pairwise ordinary least-squares linear regression was used to compare experimentally measured DVC fields across image-processing and analysis conditions, separately for annulus fibrosus fibres, cartilage endplate (CEP) and vertebral endplate (VEP) tissues. The displacement components (u, v and w) and first and third principal Green–Lagrange strains (ε₁ and ε₃) were analysed. For comparisons using an identical DVC point cloud, corresponding measurements were matched by their iDVC point identifier, n. Non-finite pairs were excluded and a linear model with an intercept was fitted to each pair of fields. The coefficient of determination was calculated as

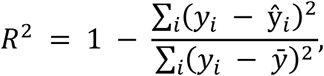

where y_i_ is the observed value, ŷ_i_ is the fitted value and ȳ is the mean observed value. R² was not calculated when fewer than three valid pairs were available or when either field had zero variance.

Regressions were performed pointwise and after regional averaging within aligned, non-overlapping cubes with side lengths of 50 or 100 voxels (81.25 or 162.5 μm, respectively). A common cube origin was defined for each tissue, and mean u, v, w, ε₁ and ε₃ values were calculated within each occupied cube before corresponding regions were matched by their three-dimensional cube indices. Pairwise R² values were displayed as regression matrices; self-comparisons were excluded from summary values. Analyses were performed using custom Python scripts with NumPy and pandas, and matrices were visualised using Matplotlib.

### Comparison of fibre-derived and regular-grid strain fields

To quantify the effect of point-cloud geometry on the resulting strain field, 1^st^ and 3^rd^ principal strain images obtained using the fibre-derived point cloud were compared with the corresponding strain images obtained using a regular Cartesian grid. The regular grid had a point spacing of 6.45 voxels, selected to approximately match the point density of the fibre-derived point cloud within the analysed volume. All strain fields were resampled onto the image voxel grid before comparison, giving aligned volumes of 199 x 249 x 219 voxels.

Similarity between corresponding strain fields was first assessed on a voxel-by-voxel basis. For each principal strain component, strain values from the regular-grid analysis were regressed against the corresponding values from the fibre-derived analysis using ordinary least-squares linear regression. The fitted slope and intercept, Pearson correlation coefficient (*r*) and coefficient of determination (*R*^2^) were calculated. Agreement in strain magnitude was additionally quantified using the root-mean-square error (RMSE) and mean absolute error (MAE) between corresponding values (Supplementary Table 2,3).

To assess whether agreement depended on spatial scale, the analysis was repeated after spatial averaging using non-overlapping cubic windows with side lengths of 10, 20, 50 and 100 image voxels. Within each window, the mean strain was calculated independently for the fibre-derived and regular-grid strain fields, and regression and error metrics were calculated between the corresponding spatially averaged values. Only complete averaging windows were included; regions at the image boundaries that did not form a complete cube were excluded. This resulted in 9,576, 1,080, 48 and 4 paired spatially averaged measurements for window sizes of 10, 20, 50 and 100 voxels, respectively. Owing to the small number of independent regions available at a 100-voxel averaging scale, these data were used only as a qualitative indication of large-scale agreement (Supplementary Table 2,3).

### Bit-depth conversion

The influence of image bit depth was assessed by comparing the original 16-bit reconstructed volumes with two 8-bit conversion strategies. For histogram-adjusted conversion, grayscale intensities were linearly rescaled between 17,000 and 34,000, based on visual inspection of the image histogram, before conversion to the 8-bit intensity range of 0–255. Intensities outside the selected range were clipped.

For full-range conversion, the complete 16-bit intensity range of 0–65,535 was linearly mapped to 0–255 without adjustment for the intensity distribution of the reconstructed images. Conversion was performed using Avizo, and identical intensity-mapping parameters were applied to each reference/deformed image pair.

DVC was performed using the same tissue masks, point clouds, correlation settings and strain-calculation parameters for the 16-bit and both 8-bit datasets. Differences in displacement and strain MAER and SDER were expressed relative to the corresponding 16-bit values.

### Strain Tensor Analysis

Radial and circumferential strains were calculated by expressing the strain tensor in a local coordinate system describing the annulus fibrosus geometry (Fig. 7civ,d). In each *x*-*y* image slice, the inner and outer boundaries of the reference-state annulus fibrosus segmentation were extracted and resampled to 100 approximately equally spaced points. A centreline and two intermediate contours halfway between the centreline and each boundary produced five approximately concentric paths. Each DVC measurement point was assigned to the nearest path in the corresponding slice. The local circumferential unit vector, ĉ, was calculated tangentially using a smoothed finite difference over five contour segments, and the in-plane radial unit vector, ř, was defined perpendicular to this tangent. The corresponding normal strains were calculated by tensor projection:

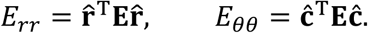

Fibre strain was calculated independently from the DVC displacement field rather than by directly projecting the calculated strain tensor (Fig. 7aiii,cii-iii). Reference-state fibre coordinates were sampled at a prescribed interval, *Δs*, and third-order polynomial space curves were fitted to the *x*, *y*, and *z* coordinates as functions of arc length, *s*.

Differentiation of these curves provided the local fibre tangent, **t**(*s*). DVC displacement vectors, **u**(*s*), were projected along the fibre:

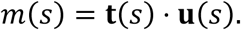

To reduce pointwise displacement noise, *m*(*s*) was fitted along each fibre using a polynomial weighted according to the iDVC objective-function minimum, *O_j_*, at each point:

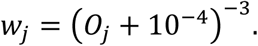

The polynomial order increased with fibre length and was calculated as 2 + round(*NΔs*/320), capped at nine and at *N* − 1, where *N* is the number of points on the fibre. The fitted displacement function, *m^~^*(*s*), was differentiated analytically, and the one-dimensional fibre-aligned Green-Lagrange strain was calculated as

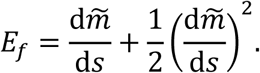

Finite-difference and unweighted polynomial estimates were also generated for comparison, while the objective-weighted estimate was used for the fibre-strain representation in Fig. 7aiii. Strains were expressed as percentages for visualisation. Analyses were implemented using the Python scripts available in the TomoSAXS repository (https://github.com/DiamondLightSource/tomosaxs).

## Data availability

The DVC, image characteristics, radial autocorrelation results, and virtual deformation images from this study are available in the Supplementary Dataset (https://doi.org/10.5522/04/33733111). The experimental deformation images used in this study are available at https://doi.org/10.5522/04/26789212, and an additional rat IVD *in situ* sCT imaging dataset suitable for running DVC is available at https://doi.org/10.5522/04/30286255.

## Code availability

Code and documentation for iDVC is available at https://tomographicimaging.github.io/iDVC/. Strain tensor and fibre based DVC analysis code is available at https://github.com/DiamondLightSource/tomosaxs.

## Acknowledgements

Laboratory space and facilities were provided by the Research Complex at Harwell. We thank S. Marussi for his support with the adaptation of the mechanical testing rigs. We acknowledge the University of Manchester at Harwell for providing the mechanical testing rig. We thank E. Newham, H.S. Gupta, J. Chen, J. Liu, S. Marathe, and Y. Zhou for their help on beamtimes. We thank the Collaborative Computational Project in Tomographic Imaging (CCPi) for supporting the development of iDVC.

## Author contributions

Conceptualization, A.L.P. and P.D.L.; methodology, A.L.P.; software, A.L.P. and B.K.B; validation, A.L.P.; formal analysis, A.L.P.; investigation, A.L.P.; resources, B.K.B. and P.D.L.; data curation, A.L.P.; writing—original draft, A.L.P., A.S. and P.D.L.; writing— review & editing, all authors; visualization, A.L.P.; supervision, P.D.L.; project administration: P.D.L.; funding acquisition, A.L.P, B.K.B., and P.D.L.

## Supplementary Information

Supplementary Information containing 3 supplementary tables, 21 supplementary figures, and 1 supplementary note. The file provides autocorrelation plots, DVC parameters, additional results plots and regression analyses.

