## Supplementary Information for "Optimising digital volume correlation across materials: a practical framework for accuracy and spatial resolution"

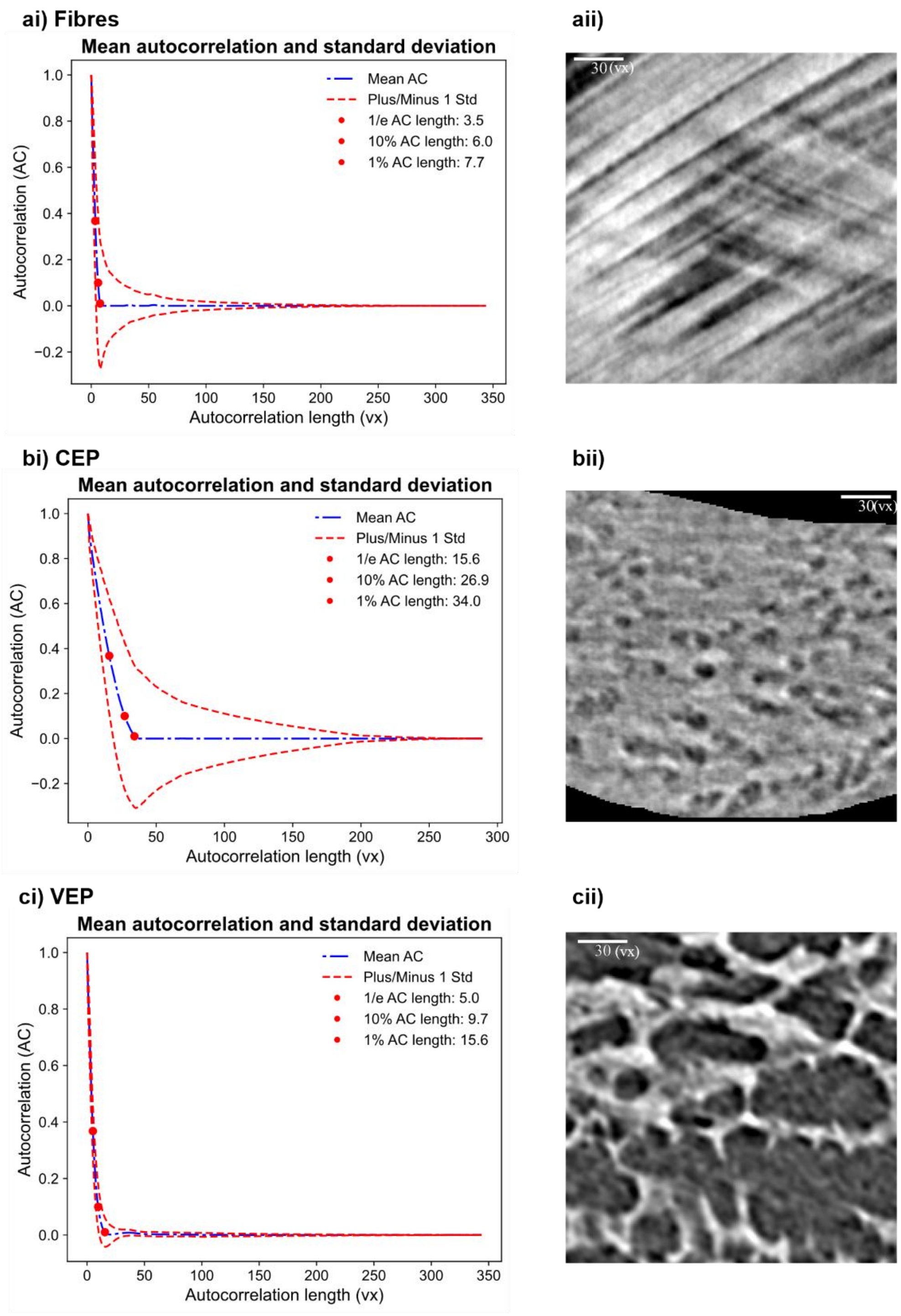


**Supplementary Figure 1 - i) Angularly averaged radial autocorrelation curve with a ±1 standard deviation ($\sigma$) envelope. The extracted 1/e, 0.1 and 0.01 correlation lengths are indicated, quantifying characteristic feature size, mesoscale, and long-range organization; ii) greyscale images of a) annulus fibrosus fibres, b) hyaline cartilage in the cartilage endplate, c) mineralised tissues (bone and cartilage) in the vertebral endplate.**

**Supplementary Table 1 – DVC parameters used for each test.**

| Test | δ/β | Tissue | Min. dist (voxels) | Subvolume size | Sampling points | Disp_max (voxels) | Strain window (points) | Number of points |
|---|---|---|---|---|---|---|---|---|
| Image filtering (virtual comp.) | ***Varied*** | Fibres | 4 | 40 | 6000 | 10 | 60 | 3387 |
| | | CEP | 5 | 40 | 6000 | 10 | 60 | 10900 |
| | | VEP | 4 | 30 | 2000 | 10 | 25 | 3700 |
| Image filtering (real strain) | ***Varied*** | Fibres | 4 | 40 | 10000 | 5 | 60 | 5000 |
| | | CEP | 5 | 40 | 6000 | 5 | 60 | 5000 |
| | | VEP | 4 | 30 | 2000 | 5 | 25 | 5000 |
| Bit-depth (virtual comp.) | 2 | Fibres | 4 | 40 | 6000 | 10 | 60 | 3387 |
| | | CEP | 5 | 40 | 6000 | 10 | 60 | 10900 |
| | | VEP | 4 | 30 | 2000 | 10 | 25 | 3700 |
| Bit-depth (real strain) | 2 | Fibres | 4 | 40 | 10000 | 5 | 60 | 5000 |
| | | CEP | 5 | 40 | 6000 | 5 | 60 | 5000 |
| | | VEP | 4 | 30 | 2000 | 5 | 25 | 5000 |
| Point density (virtual comp.) | 2 | Fibres | ***Varied*** | 40 | 6000 | 10 | 50 | ***Varied*** |
| | | CEP | ***Varied*** | 40 | 6000 | 10 | 50 | ***Varied*** |
| | | VEP | ***Varied*** | 30 | 2000 | 10 | 50 | ***Varied*** |
| Point density (real strain) | 2 | Fibres | ***Varied*** | 40 | 6000 | 5 | 50 | ***Varied*** |
| | | CEP | ***Varied*** | 40 | 6000 | 5 | 50 | ***Varied*** |
| | | VEP | ***Varied*** | 30 | 2000 | 5 | 50 | ***Varied*** |
| Sub-volume size (virtual comp.) | 2 | Fibres | 4 | ***Varied*** | 6000 | 10 | 75 | 3387 |
| | | CEP | 5 | ***Varied*** | 6000 | 10 | 75 | 10900 |
| | | VEP | 4 | ***Varied*** | 2000 | 10 | 75 | 3700 |
| Sub-volume size (real strain) | 2 | Fibres | 4 | ***Varied*** | 6000 | 5 | 50 | 171333 |
| | | CEP | 5 | ***Varied*** | 6000 | 5 | 50 | 16174 |
| | | VEP | 4 | ***Varied*** | 2000 | 5 | 50 | 56364 |
| Strain window (virtual comp.) | 2 | Fibres | 4 | 40 | 6000 | 10 | ***Varied*** | 3387 |
| | | CEP | 4 | 40 | 6000 | 10 | ***Varied*** | 19356 |
| | | VEP | 4 | 30 | 2000 | 10 | ***Varied*** | 3700 |
| Strain window (real strain) | 2 | Fibres | 4 | 40 | 6000 | 5 | ***Varied*** | 54668 |
| | | CEP | 4 | 40 | 6000 | 5 | ***Varied*** | 30908 |
| | | VEP | 4 | 30 | 2000 | 5 | ***Varied*** | 56354 |

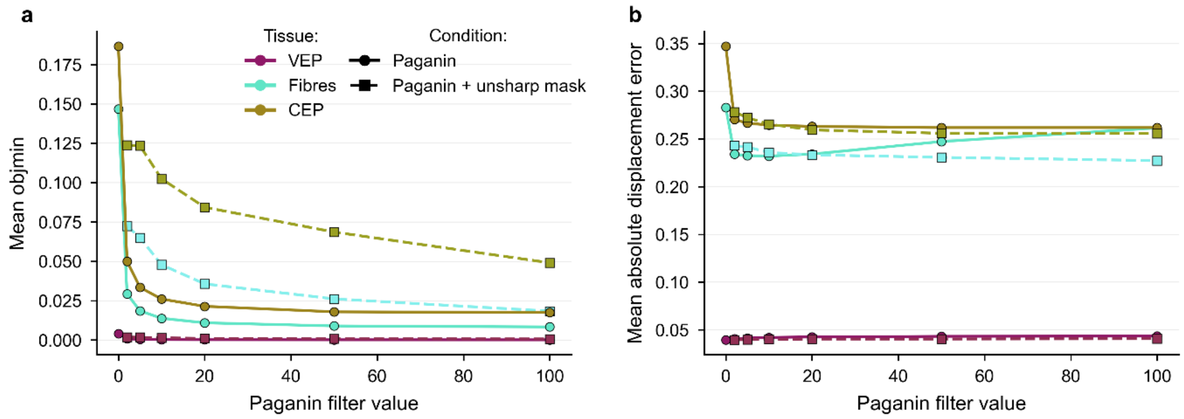


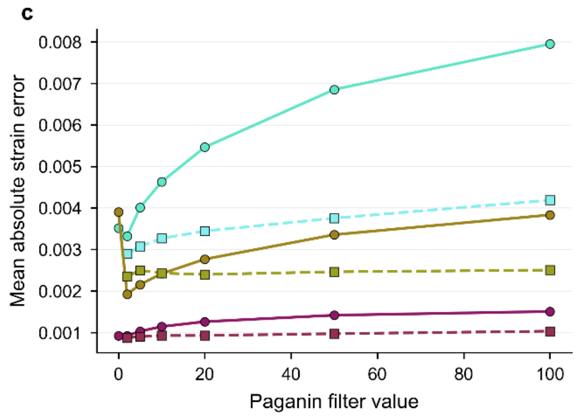


**d**

| Tissue | Lowest strain MAER | Lowest displacement MAER |
|---|---|---|
| VEP | 0.000875<br>Paganin 2<br>+ unsharp mask | 0.0394<br>Paganin 0 |
| Fibres | 0.0029<br>Paganin 2<br>+ unsharp mask | 0.227<br>Paganin 100<br>+ unsharp mask |
| CEP | 0.00192<br>Paganin 2 | 0.256<br>Paganin 100<br>+ unsharp mask |

**Supplementary Figure 2 – Virtual compression analysis on the impact of image filtering on a) mean objective minimum values, b) mean absolute displacement error, and c) mean absolute strain error; d) the filter settings that gave the lowest strain and displacement errors for each tissue type.**

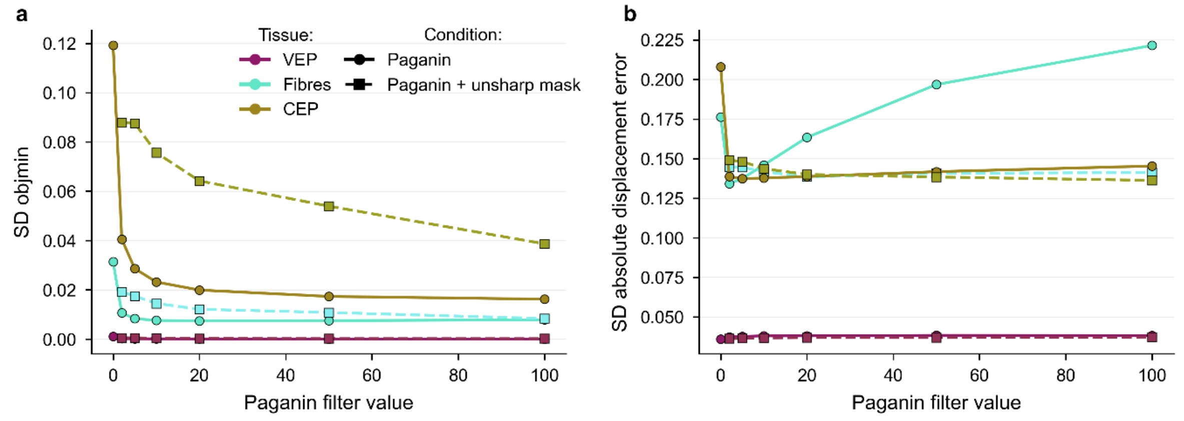


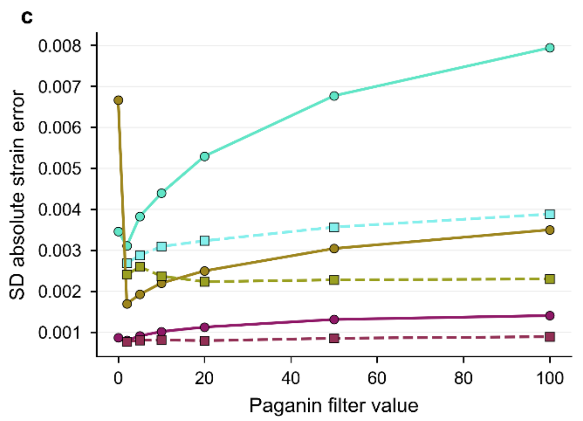


d

| Tissue | Lowest strain SDER | Lowest displacement SDER |
|---|---|---|
| VEP | 0.000766<br>Paganin 2<br>+ unsharp mask | 0.0361<br>Paganin 0 |
| Fibres | 0.00269<br>Paganin 2<br>+ unsharp mask | 0.134<br>Paganin 2 |
| CEP | 0.00169<br>Paganin 2 | 0.136<br>Paganin 100<br>+ unsharp mask |

**Supplementary Figure 3 - Virtual compression analysis on the impact of image filtering on a) standard deviation of objective minimum values, b) standard deviation of absolute displacement error, and c) standard deviation of absolute strain error; d) the filter settings that gave the lowest strain and displacement errors for each tissue type.**

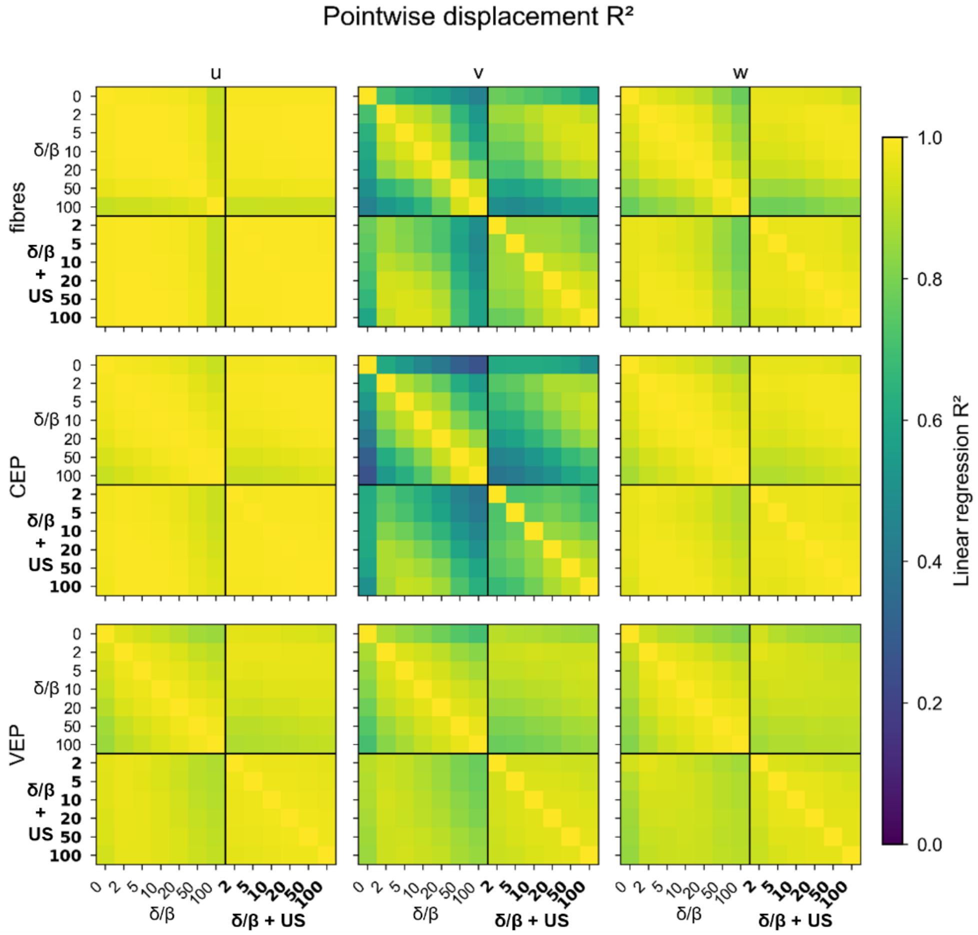


**Supplementary Figure 4 – Pointwise linear regression analysis of displacement components across different filtering parameters for each tissue.**

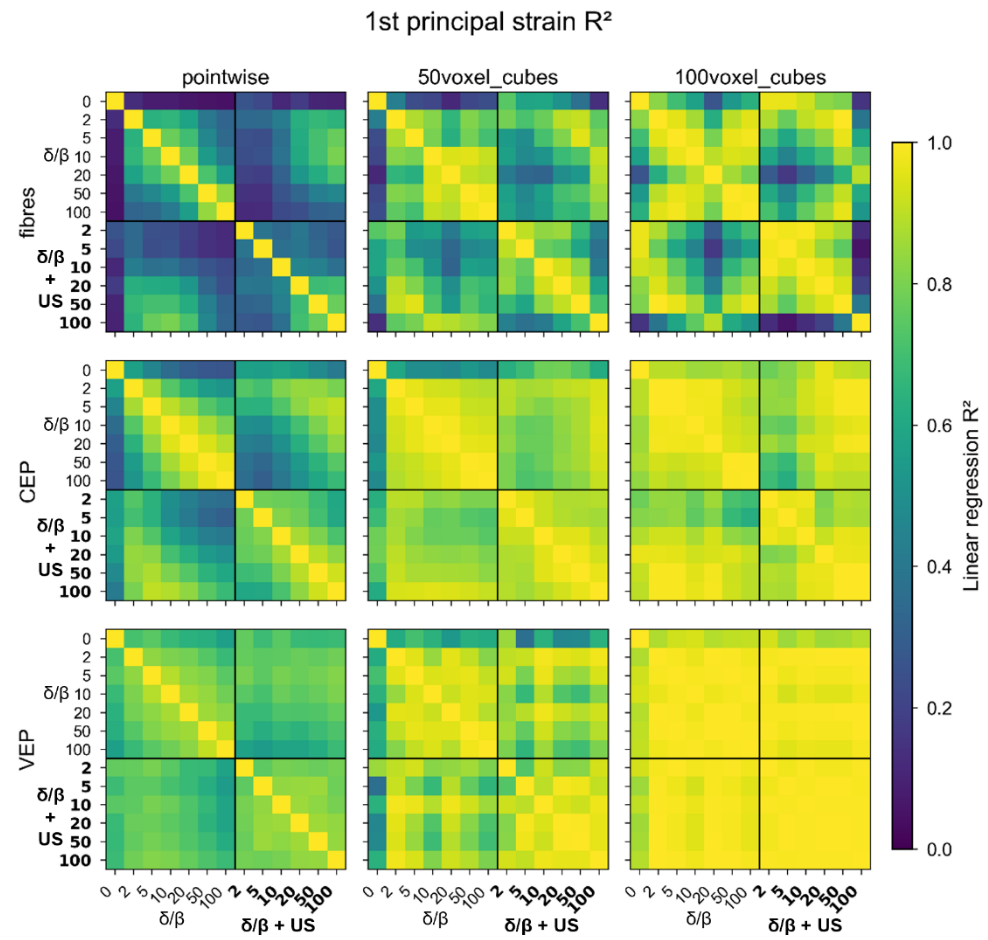


**Supplementary Figure 5 – Linear regression analysis of 1st principal strain fields across different filtering parameters and spatial averaging length scales for each tissue.**

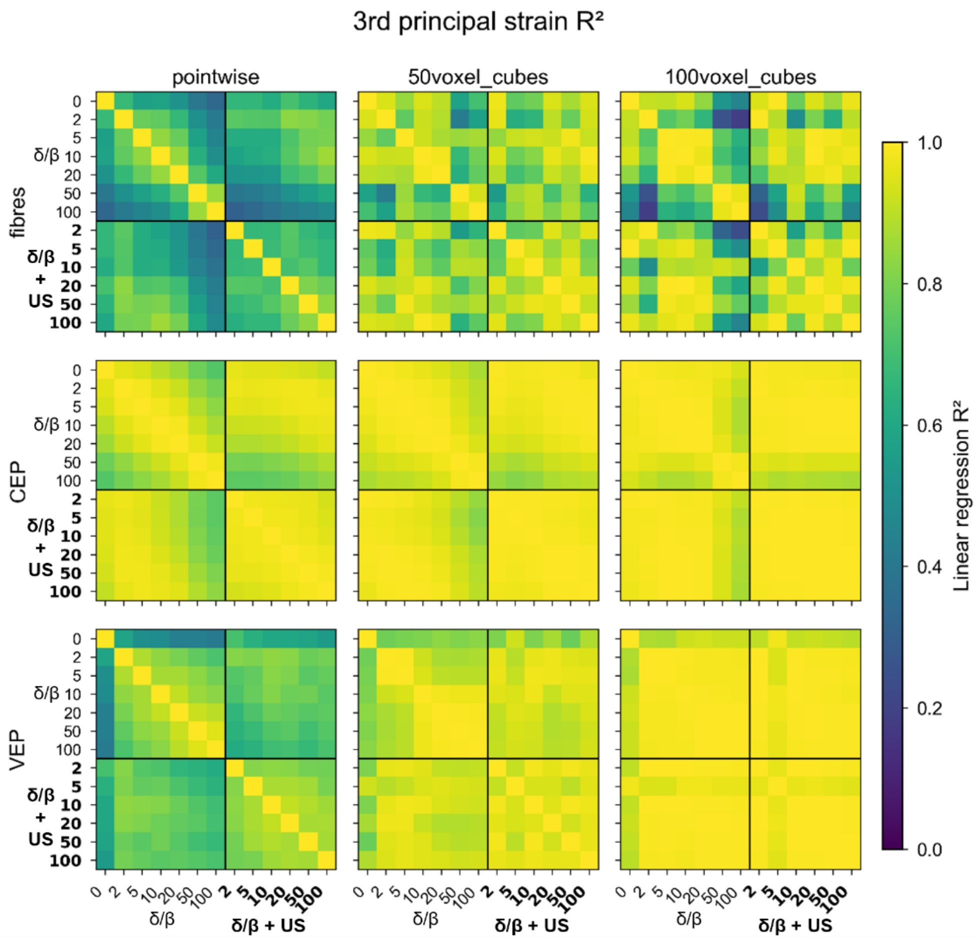


**Supplementary Figure 6 - Linear regression analysis of 3rd principal strain fields across different filtering parameters and spatial averaging length scales for each tissue.**

### Supplementary Note 1 – Summary of linear regression across different filtering parameters

Values use unique off-diagonal filter pairs only; diagonal self-comparisons and mirrored A–B/B–A duplicates are excluded.

#### *3rd principal strain*

*Pointwise*:

AF fibres: median $R^2$ = 0.66 (IQR 0.54–0.74; range 0.31–0.86; 49/78 [63%] filter pairs below $R^2$ = 0.70).

CEP cartilage: median $R^2$ = 0.95 (IQR 0.88–0.97; range 0.73–0.99; 0/78 [0%] filter pairs below $R^2$ = 0.70).

VEP: median $R^2$ = 0.78 (IQR 0.70–0.83; range 0.41–0.95; 19/78 [24%] filter pairs below $R^2$ = 0.70).

*50-voxel-cube averaged*:

AF fibres: median $R^2$ = 0.87 (IQR 0.77–0.93; range 0.43–0.99; 10/78 [13%] filter pairs below $R^2$ = 0.70).

CEP cartilage: median $R^2$ = 0.98 (IQR 0.94–0.99; range 0.82–1.00; 0/78 [0%] filter pairs below $R^2$ = 0.70).

VEP: median $R^2$ = 0.94 (IQR 0.90–0.96; range 0.78–1.00; 0/78 [0%] filter pairs below $R^2$ = 0.70).

*100-voxel-cube averaged*:

AF fibres: median $R^2$ = 0.89 (IQR 0.70–0.96; range 0.19–1.00; 20/78 [26%] filter pairs below $R^2$ = 0.70).

CEP cartilage: median $R^2$ = 0.99 (IQR 0.96–1.00; range 0.87–1.00; 0/78 [0%] filter pairs below $R^2$ = 0.70).

VEP: median $R^2$ = 0.99 (IQR 0.97–1.00; range 0.87–1.00; 0/78 [0%] filter pairs below $R^2$ = 0.70).

***Pointwise strain agreement***

AF fibres, 1st principal strain: median $R^2$ = 0.37 (IQR 0.24–0.61; range 0.05–0.84; 68/78 [87%] filter pairs below $R^2$ = 0.70).

AF fibres, 3rd principal strain: median $R^2$ = 0.66 (IQR 0.54–0.74; range 0.31–0.86; 49/78 [63%] filter pairs below $R^2$ = 0.70).

CEP cartilage, 1st principal strain: median $R^2$ = 0.69 (IQR 0.52–0.81; range 0.26–0.97; 40/78 [51%] filter pairs below $R^2$ = 0.70).

CEP cartilage, 3rd principal strain: median $R^2$ = 0.95 (IQR 0.88–0.97; range 0.73–0.99; 0/78 [0%] filter pairs below $R^2$ = 0.70).

VEP, 1st principal strain: median $R^2$ = 0.76 (IQR 0.68–0.80; range 0.55–0.91; 22/78 [28%] filter pairs below $R^2$ = 0.70).

VEP, 3rd principal strain: median $R^2$ = 0.78 (IQR 0.70–0.83; range 0.41–0.95; 19/78 [24%] filter pairs below $R^2$ = 0.70).

***Pointwise displacement agreement***

AF fibres, u displacement: median $R^2$ = 0.99 (IQR 0.98–1.00; range 0.91–1.00; 0/78 [0%] filter pairs below $R^2$ = 0.70).

AF fibres, v displacement: median $R^2$ = 0.81 (IQR 0.66–0.90; range 0.44–0.95; 21/78 [27%] filter pairs below $R^2$ = 0.70).

AF fibres, w displacement: median $R^2$ = 0.96 (IQR 0.91–0.97; range 0.77–0.99; 0/78 [0%] filter pairs below $R^2$ = 0.70).

CEP cartilage, u displacement: median $R^2$ = 0.99 (IQR 0.97–0.99; range 0.92–1.00; 0/78 [0%] filter pairs below $R^2$ = 0.70).

CEP cartilage, v displacement: median $R^2$ = 0.73 (IQR 0.61–0.84; range 0.25–0.96; 33/78 [42%] filter pairs below $R^2$ = 0.70).

CEP cartilage, w displacement: median $R^2$ = 0.97 (IQR 0.95–0.98; range 0.87–0.99; 0/78 [0%] filter pairs below $R^2$ = 0.70).

VEP, u displacement: median $R^2$ = 0.96 (IQR 0.93–0.97; range 0.85–0.98; 0/78 [0%] filter pairs below $R^2$ = 0.70).

VEP, v displacement: median $R^2$ = 0.91 (IQR 0.85–0.93; range 0.71–0.97; 0/78 [0%] filter pairs below $R^2$ = 0.70).

VEP, w displacement: median $R^2$ = 0.92 (IQR 0.90–0.95; range 0.82–0.98; 0/78 [0%] filter pairs below $R^2$ = 0.70).

***Effect of spatial averaging on strain agreement***

CEP cartilage, 1st principal strain: median $R^2$ 0.69 pointwise -> 0.87 at 50 voxels -> 0.92 at 100 voxels

CEP cartilage, 3rd principal strain: median $R^2$ 0.95 pointwise -> 0.98 at 50 voxels -> 0.99 at 100 voxels

VEP, 1st principal strain: median $R^2$ 0.76 pointwise -> 0.89 at 50 voxels -> 0.99 at 100 voxels

VEP, 3rd principal strain: median $R^2$ 0.78 pointwise -> 0.94 at 50 voxels -> 0.99 at 100 voxels

AF fibres, 1st principal strain: median $R^2$ 0.37 pointwise -> 0.70 at 50 voxels -> 0.80 at 100 voxels

AF fibres, 3rd principal strain: median $R^2$ 0.66 pointwise -> 0.87 at 50 voxels -> 0.89 at 100 voxels

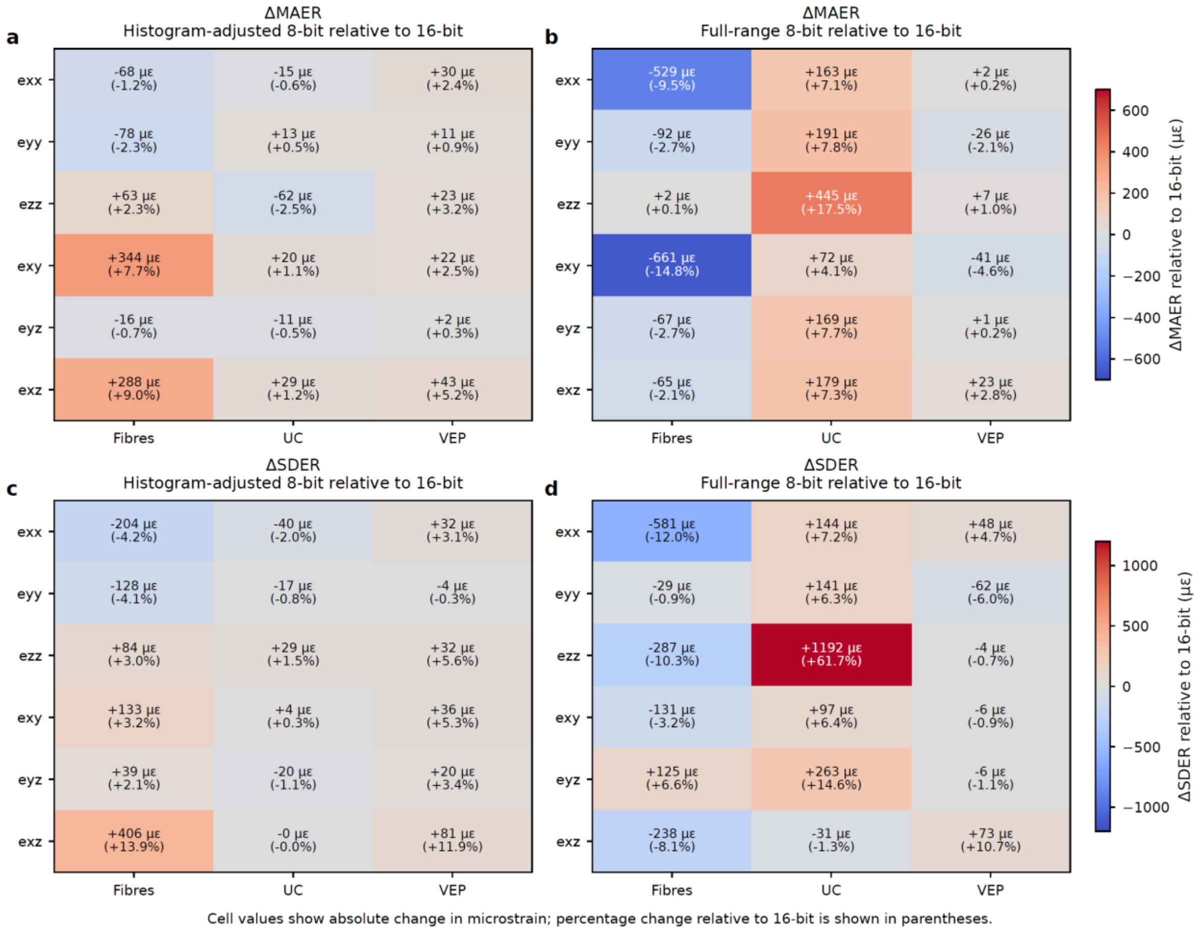


**Supplementary Figure 7 – Change in strain mean absolute error between 16-bit and a) histogram adjusted 8-bit, b) full-range 8-bit. Change in strain standard deviation of absolute error between 16-bit and c) histogram adjusted 8-bit, d) full-range 8-bit.**

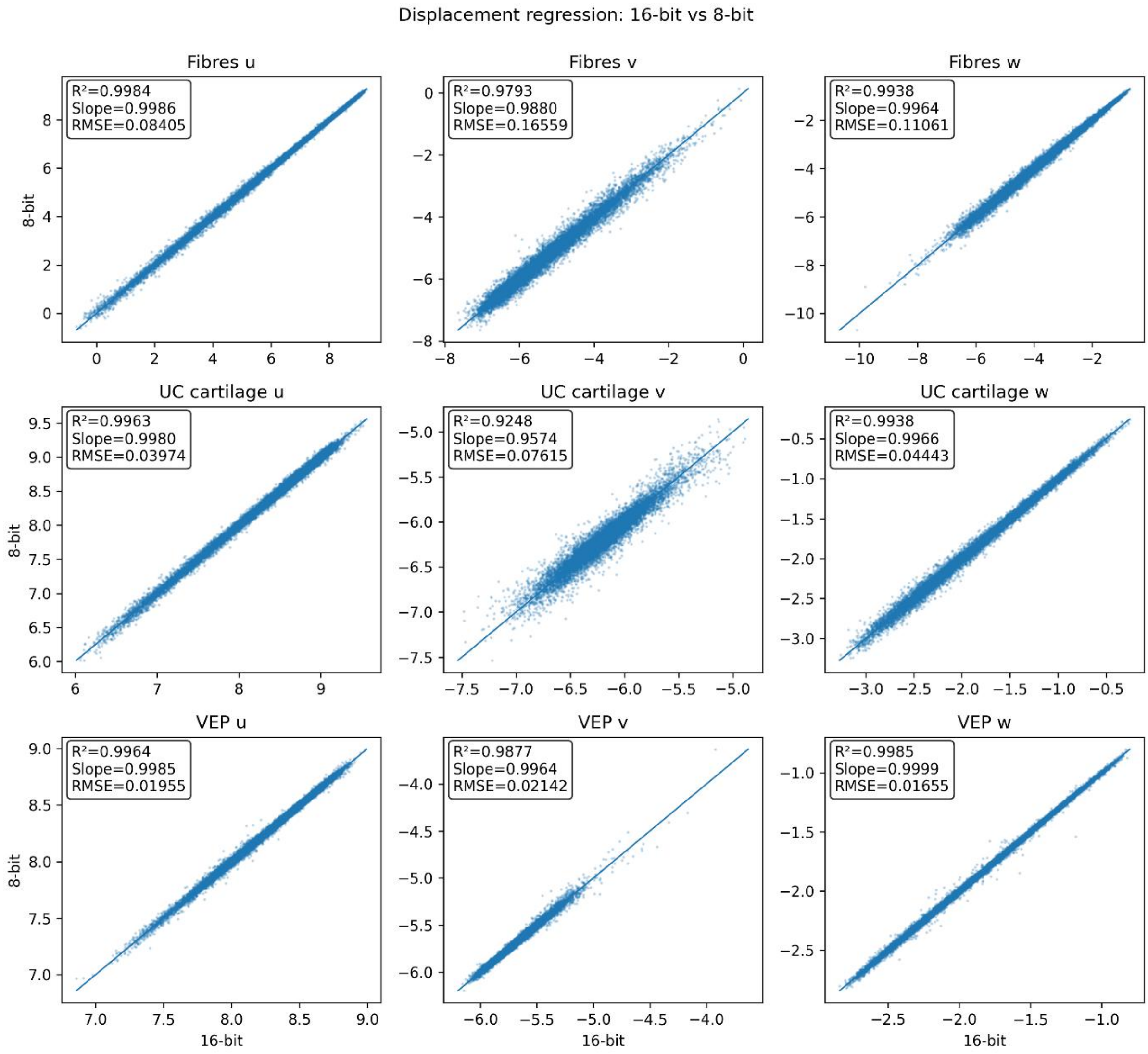


**Supplementary Figure 8 - 16-bit vs histogram adjusted 8-bit displacement component linear regression analysis**

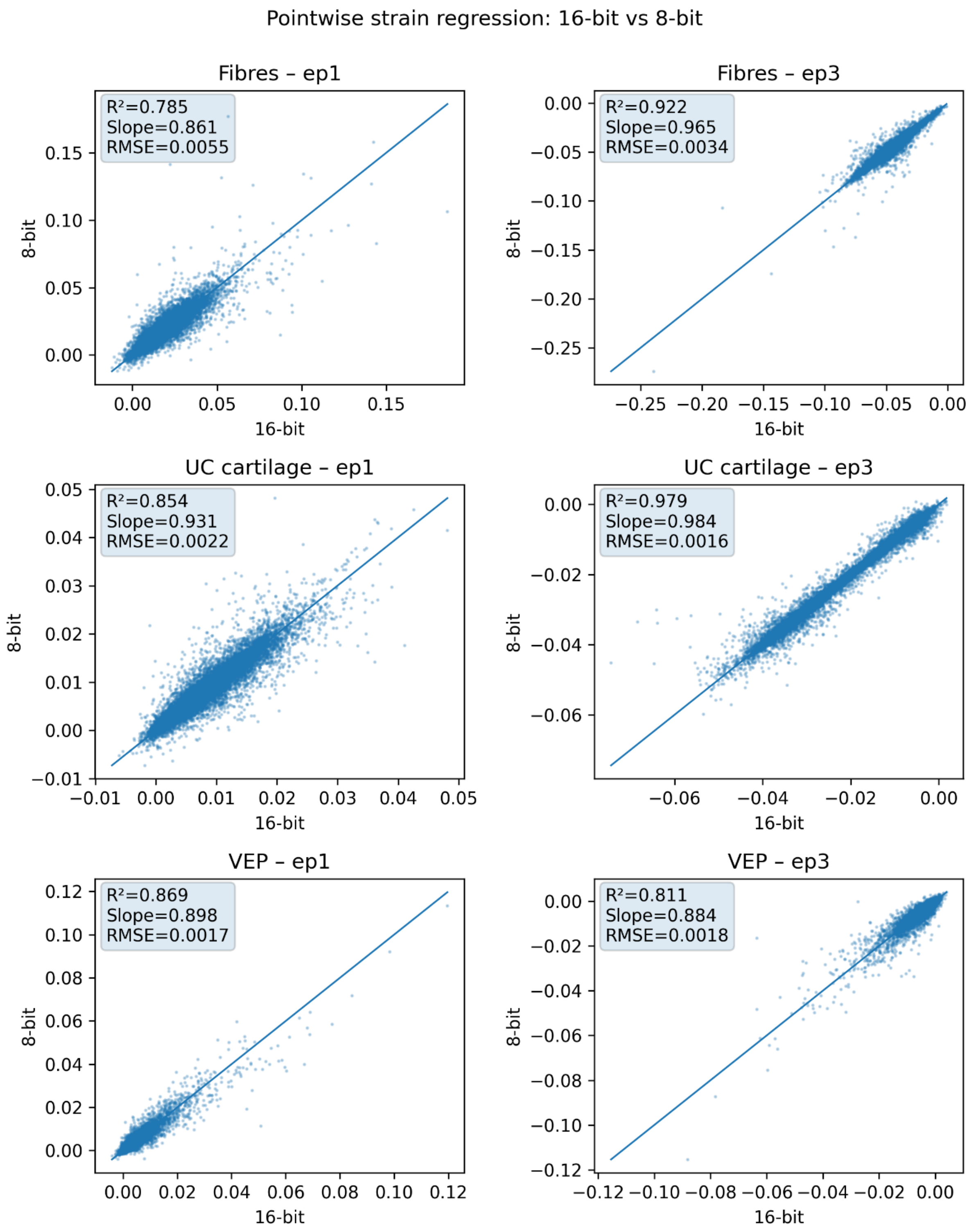


**Supplementary Figure 9 - 16-bit vs histogram adjusted 8-bit principal strain linear regression analysis.**

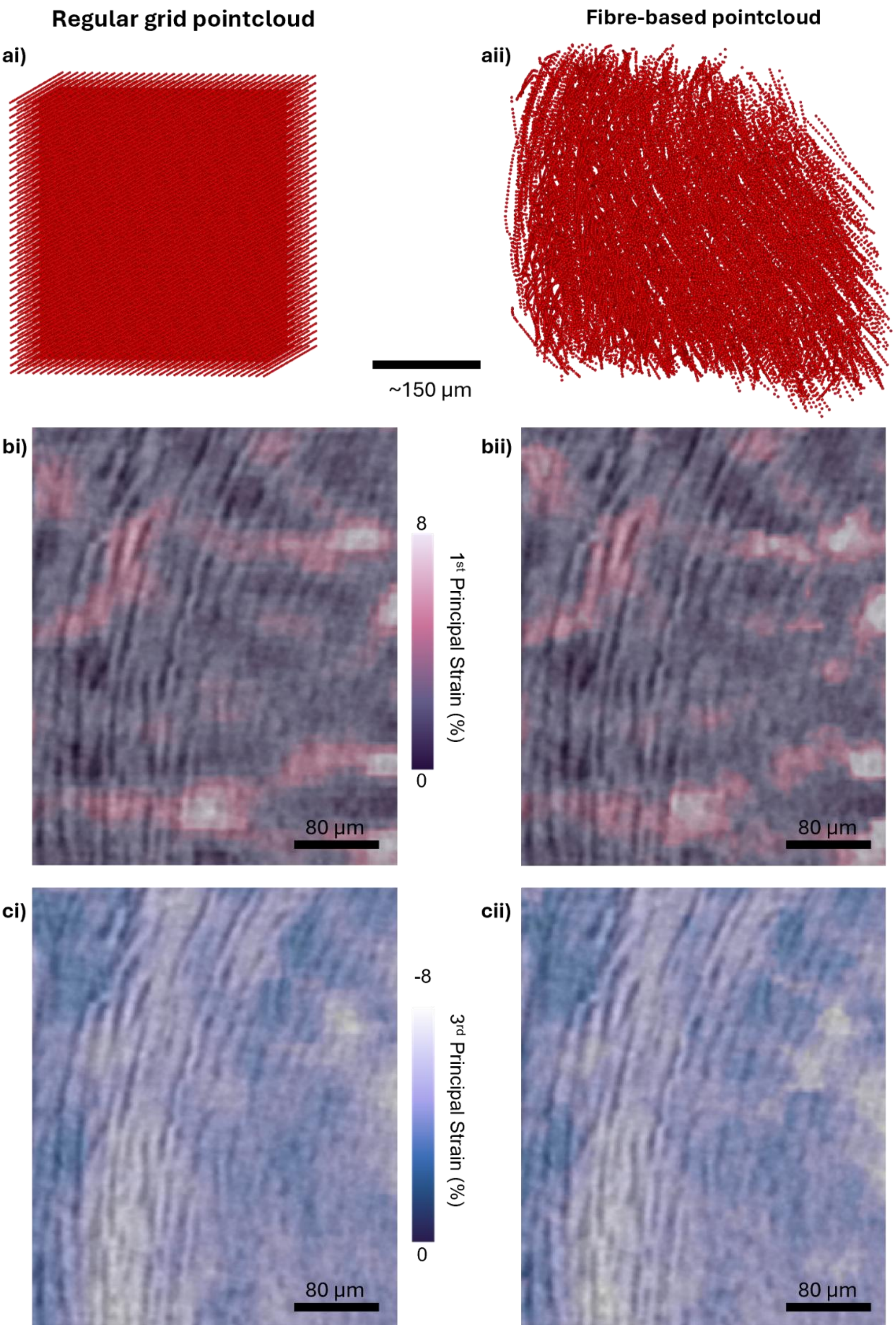


**Supplementary Figure 10 - Regular grid vs fibre-based point cloud. a) point cloud 3D rendering, b) 1st principal strain, c) 3rd principal strain for i) a regular grid point cloud and ii) a fibre-based point cloud.**

**Supplementary Table 2 – Regression analysis of 1st principal strain values obtained from a regular grid point cloud vs fibre-based point cloud.**

| Scale | Voxels/Regions | Slope | Intercept | r | $R^2$ | RMSE | MAE |
|---|---|---|---|---|---|---|---|
| Voxel-wise | 10,851,669 | 0.810 | 0.00354 | 0.827 | **0.684** | 0.01060 | 0.00674 |
| 10 voxel | 9,576 | 1.027 | -0.00126 | 0.988 | **0.976** | 0.00122 | 0.00099 |
| 20 voxel | 1,080 | 1.012 | -0.00104 | 0.996 | **0.992** | 0.00095 | 0.00084 |
| 50 voxel | 48 | 0.991 | -0.00071 | 0.999 | **0.998** | 0.00093 | 0.00090 |
| 100 voxel | 4 | 0.743 | 0.00354 | 0.969 | **0.940** | 0.00098 | 0.00098 |

**Supplementary Table 3 - Regression analysis of 3rd principal strain values obtained from a regular grid point cloud vs fibre-based point cloud.**

| Scale | Voxels/Regions | Slope | Intercept | r | $R^2$ | RMSE | MAE |
|---|---|---|---|---|---|---|---|
| Voxel-wise | 10,851,669 | 0.875 | -0.00476 | 0.881 | **0.776** | 0.00672 | 0.00411 |
| 10 voxel | 9,576 | 1.018 | 0.00128 | 0.998 | **0.995** | 0.00078 | 0.00064 |
| 20 voxel | 1,080 | 1.012 | 0.00110 | 1.000 | **0.999** | 0.00064 | 0.00059 |
| 50 voxel | 48 | 1.012 | 0.00109 | 1.000 | **0.9998** | 0.00063 | 0.00062 |
| 100 voxel | 4 | 0.522 | -0.01657 | 0.992 | **0.984** | 0.00068 | 0.00068 |

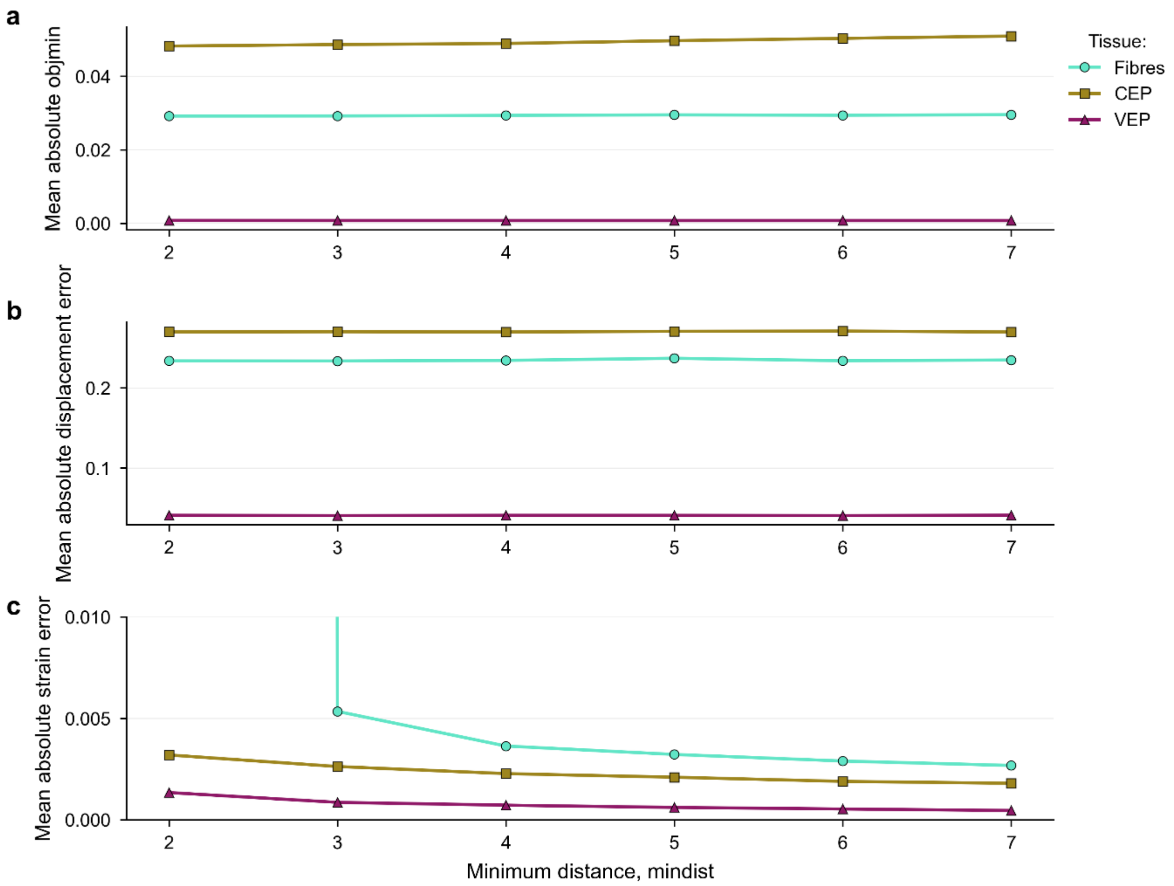


d

| Tissue | Lowest objmin MAER | Lowest displacement MAER | Lowest strain MAER |
|---|---|---|---|
| Fibres | 0.0291<br>mindist 2 | 0.233<br>mindist 3 | 0.00268<br>mindist 7 |
| CEP | 0.0482<br>mindist 2 | 0.27<br>mindist 7 | 0.0018<br>mindist 7 |
| VEP | 0.000742<br>mindist 7 | 0.0405<br>mindist 3 | 0.00046<br>mindist 7 |

**Supplementary Figure 11 - Virtual compression analysis on the impact of point spacing (minimum distance) on a) mean objective minimum values, b) mean absolute displacement error, and c) mean absolute strain error; d) the point spacings that gave the lowest strain and displacement errors for each tissue type.**

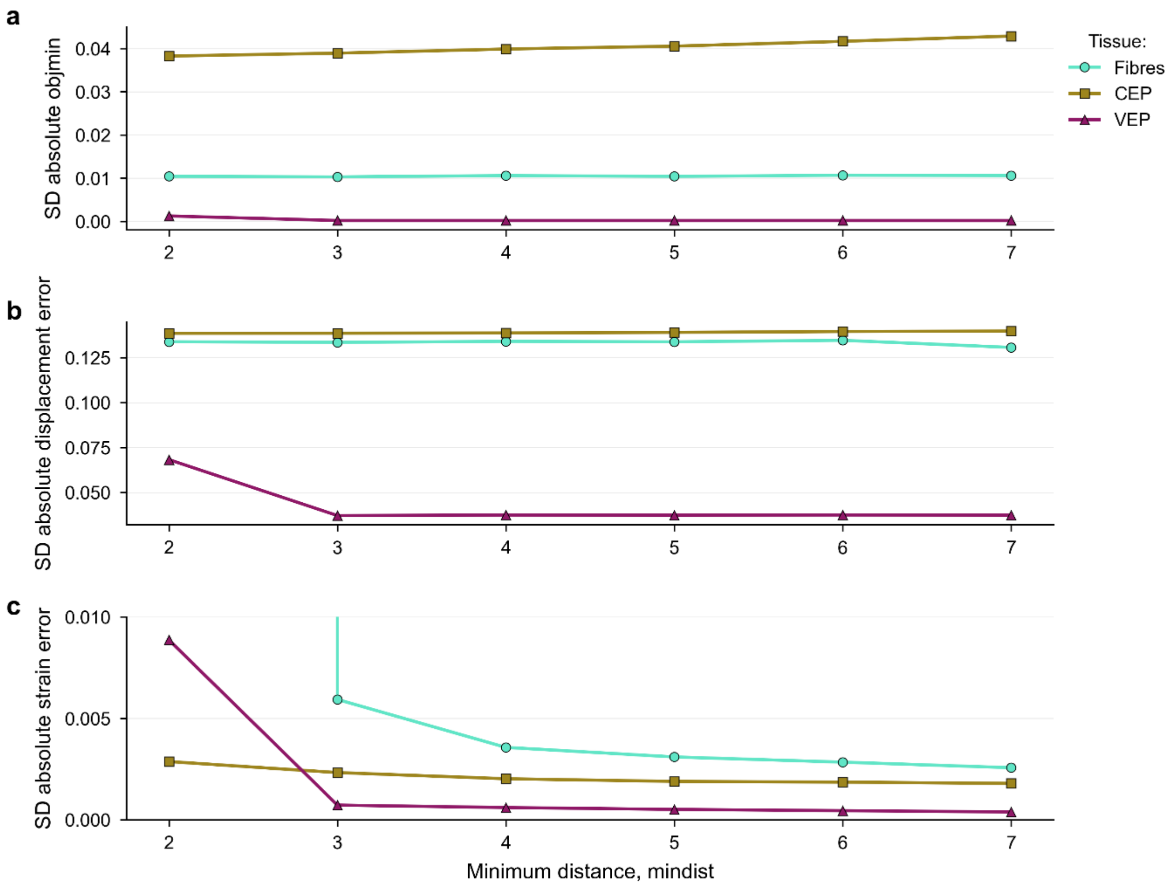


**d**

| Tissue | Lowest objmin SDER | Lowest displacement SDER | Lowest strain SDER |
|---|---|---|---|
| Fibres | 0.0103<br>mindist 3 | 0.131<br>mindist 7 | 0.00257<br>mindist 7 |
| CEP | 0.0383<br>mindist 2 | 0.138<br>mindist 2 | 0.00179<br>mindist 7 |
| VEP | 0.000212<br>mindist 7 | 0.0372<br>mindist 3 | 0.000386<br>mindist 7 |

**Supplementary Figure 12 - Virtual compression analysis on the impact of point spacing (minimum distance) on a) standard deviation (SD) objective minimum values, b) standard deviation absolute displacement error, and c) standard deviation absolute strain error; d) the point spacings that gave the lowest strain and displacement standard deviation of error for each tissue type.**

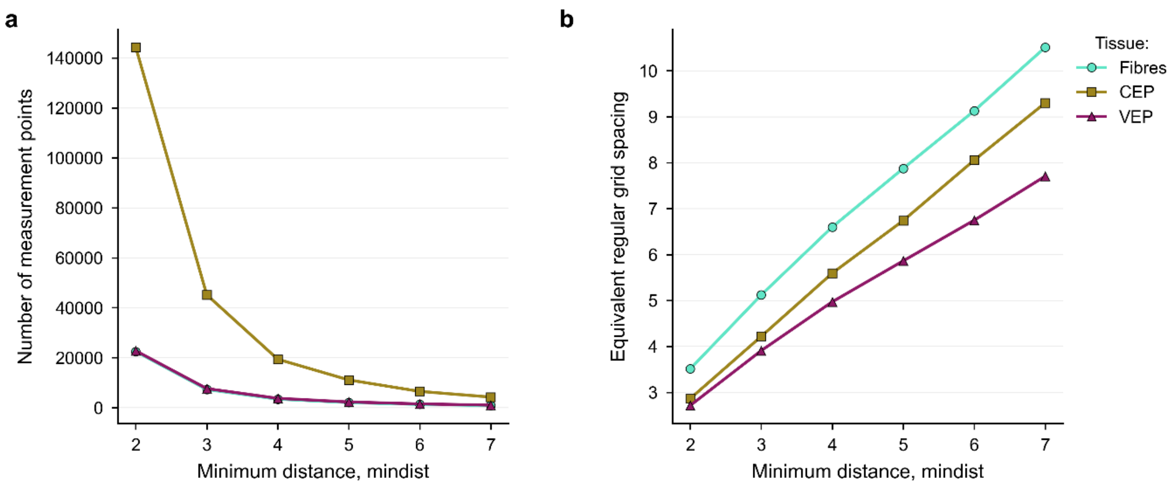


**Supplementary Figure 13 – a) number of measurement points generated for fibres, CEP, and VEP using different prescribed minimum distances (voxels) between points; b) the equivalent regular cartesian grid spacing (voxels) for point clouds generated with different minimum distances.**

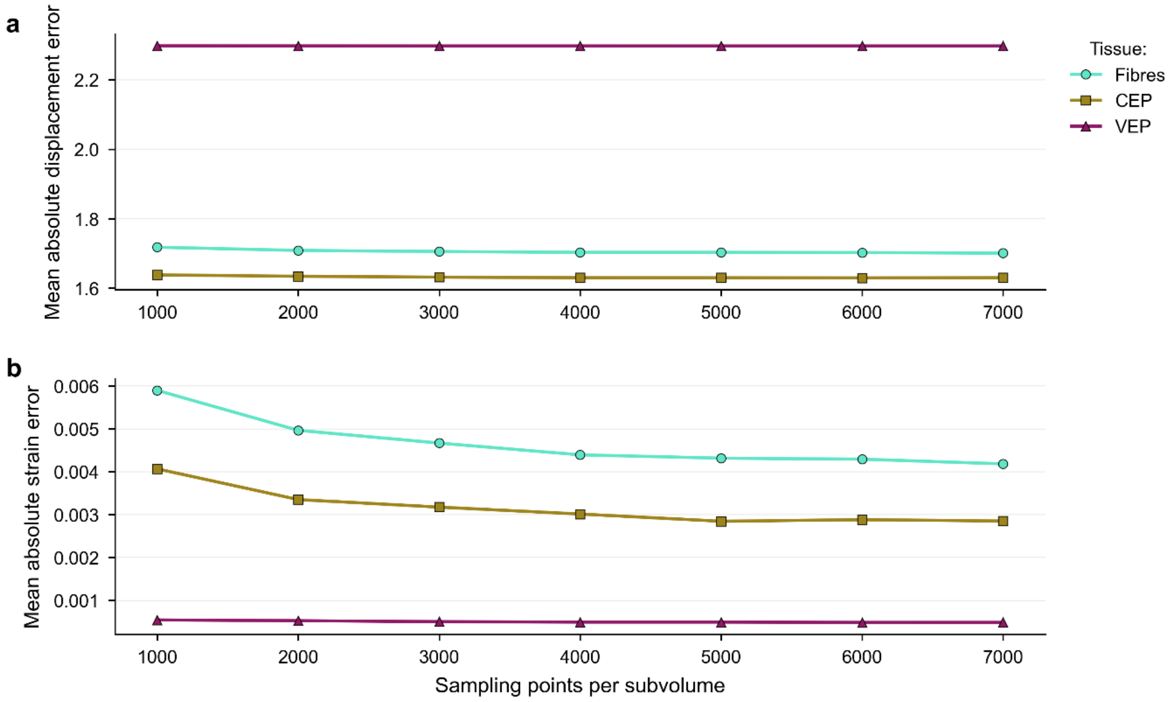


**Supplementary Figure 14 - Mean absolute error of a) displacement and b) strain with increasing sampling points per subvolume.**

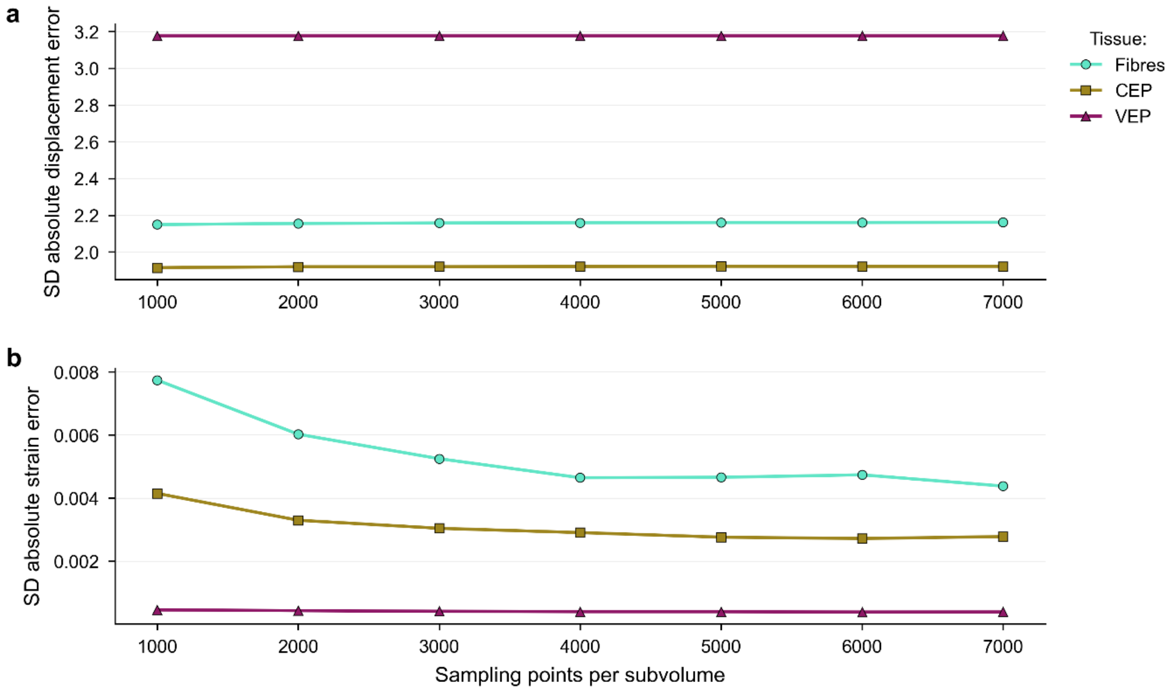


**Supplementary Figure 15 – Standard deviation of absolute error of a) displacement and b) strain with increasing sampling points per subvolume.**

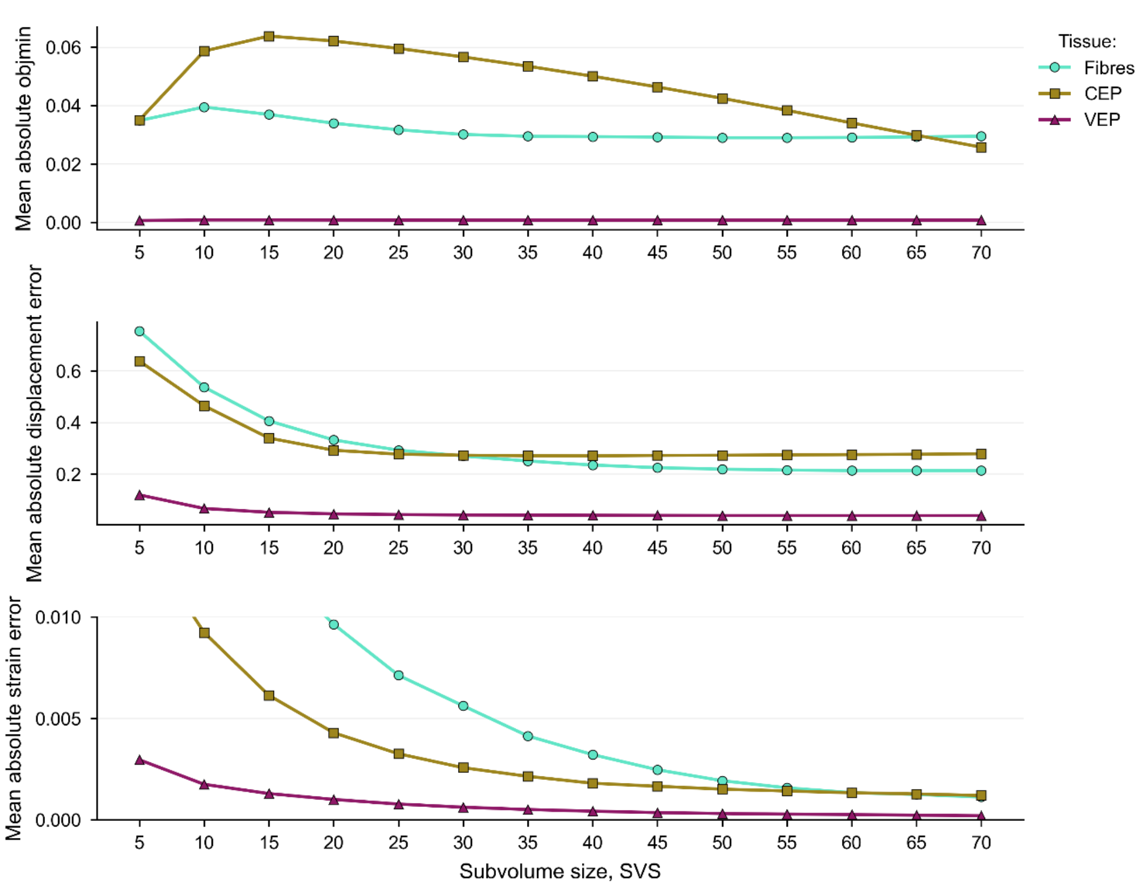


d

| Tissue | Lowest objmin MAER | Lowest displacement MAER | Lowest strain MAER |
|---|---|---|---|
| Fibres | 0.029<br>SVS 55 | 0.213<br>SVS 65 | 0.00112<br>SVS 70 |
| CEP | 0.0257<br>SVS 70 | 0.27<br>SVS 40 | 0.0012<br>SVS 70 |
| VEP | 0.000601<br>SVS 5 | 0.0383<br>SVS 65 | 0.000202<br>SVS 70 |

**Supplementary Figure 16 – Virtual compression analysis on the impact of spherical subvolume diameter (SVS) on a) mean objective minimum values, b) mean absolute displacement error, and c) mean absolute strain error; d) the subvolume diameters that gave the lowest strain and displacement errors for each tissue type.**

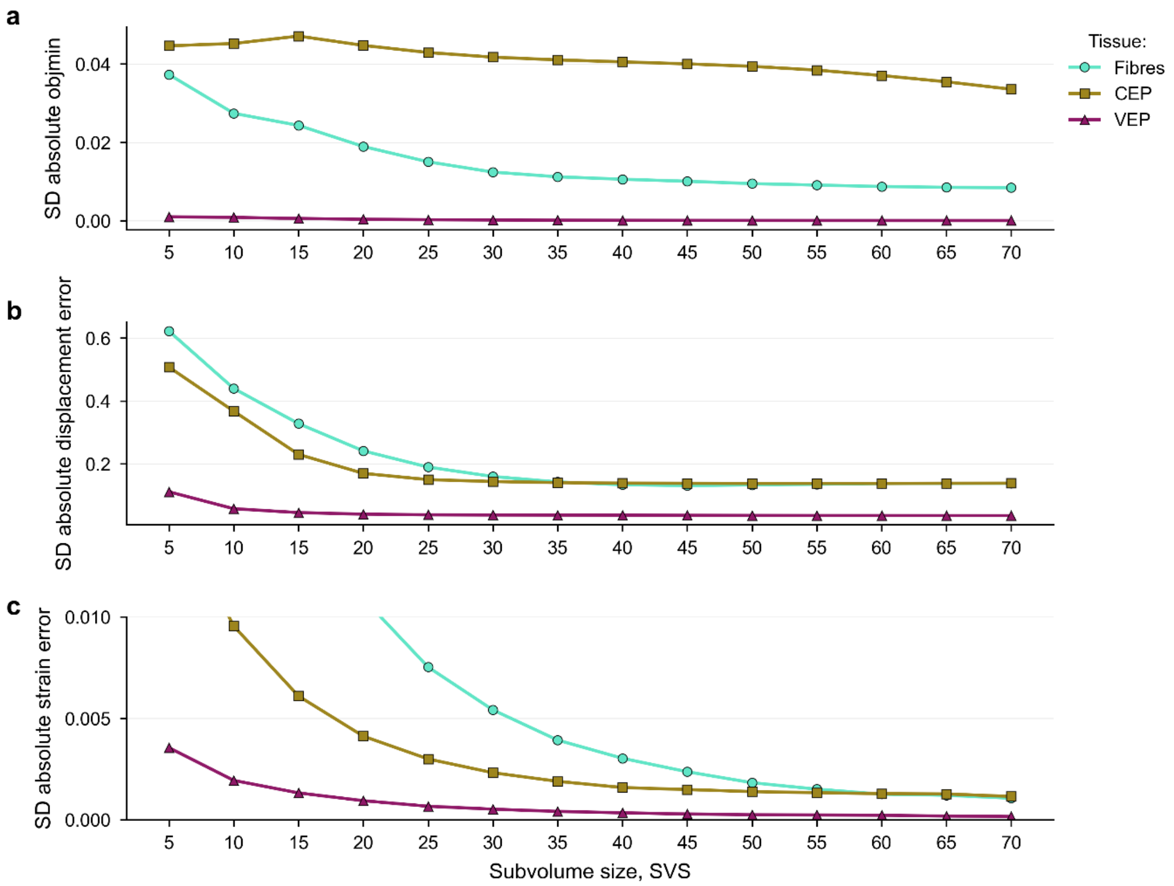


**d**

| Tissue | Lowest objmin SDER | Lowest displacement SDER | Lowest strain SDER |
|---|---|---|---|
| Fibres | 0.00845<br>SVS 70 | 0.131<br>SVS 45 | 0.00107<br>SVS 70 |
| CEP | 0.0336<br>SVS 70 | 0.138<br>SVS 60 | 0.00115<br>SVS 70 |
| VEP | 8.25e-05<br>SVS 70 | 0.0356<br>SVS 70 | 0.000161<br>SVS 70 |

**Supplementary Figure 17 – Virtual compression analysis on the impact of spherical subvolume diameter (SVS) on a) standard deviation objective minimum values, b) standard deviation absolute displacement error, and c) standard deviation absolute strain error; d) the subvolume diameters that gave the lowest strain and displacement standard deviation of errors for each tissue type.**

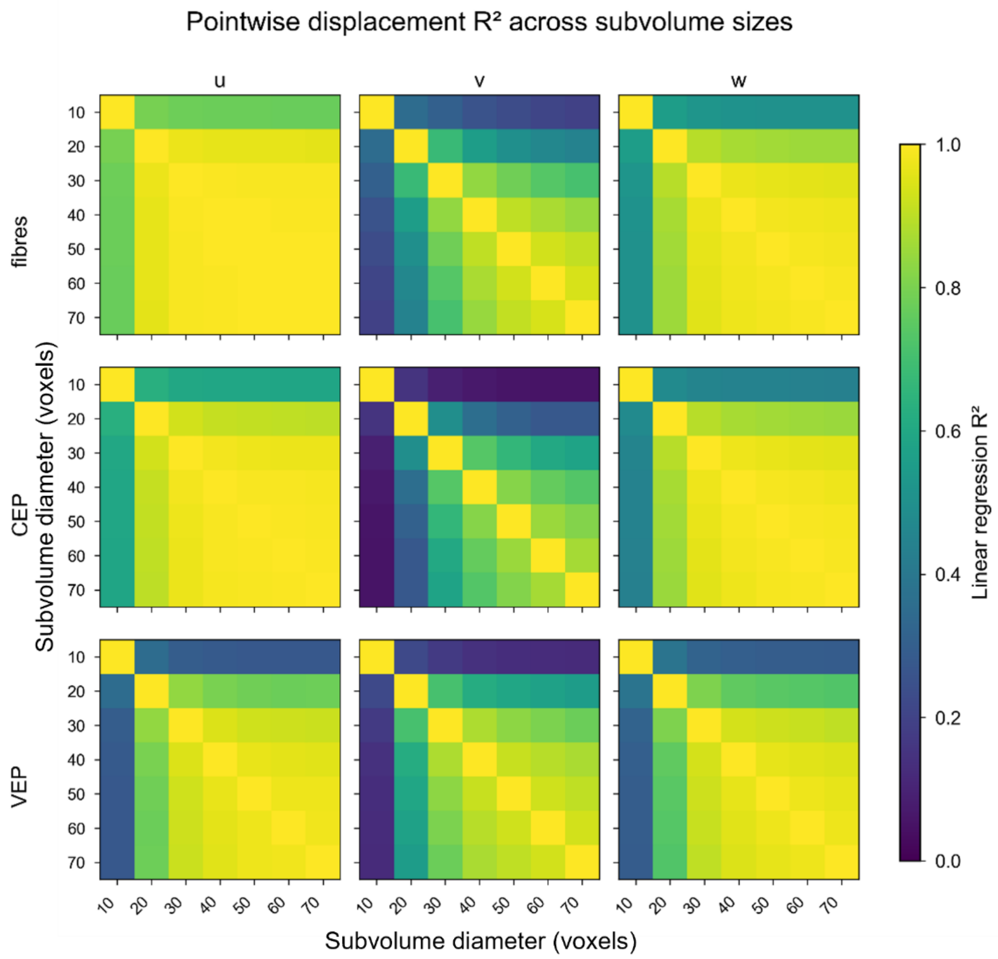


**Supplementary Figure 18 - Pointwise linear regression analysis of displacement components across different subvolume sizes for each tissue.**

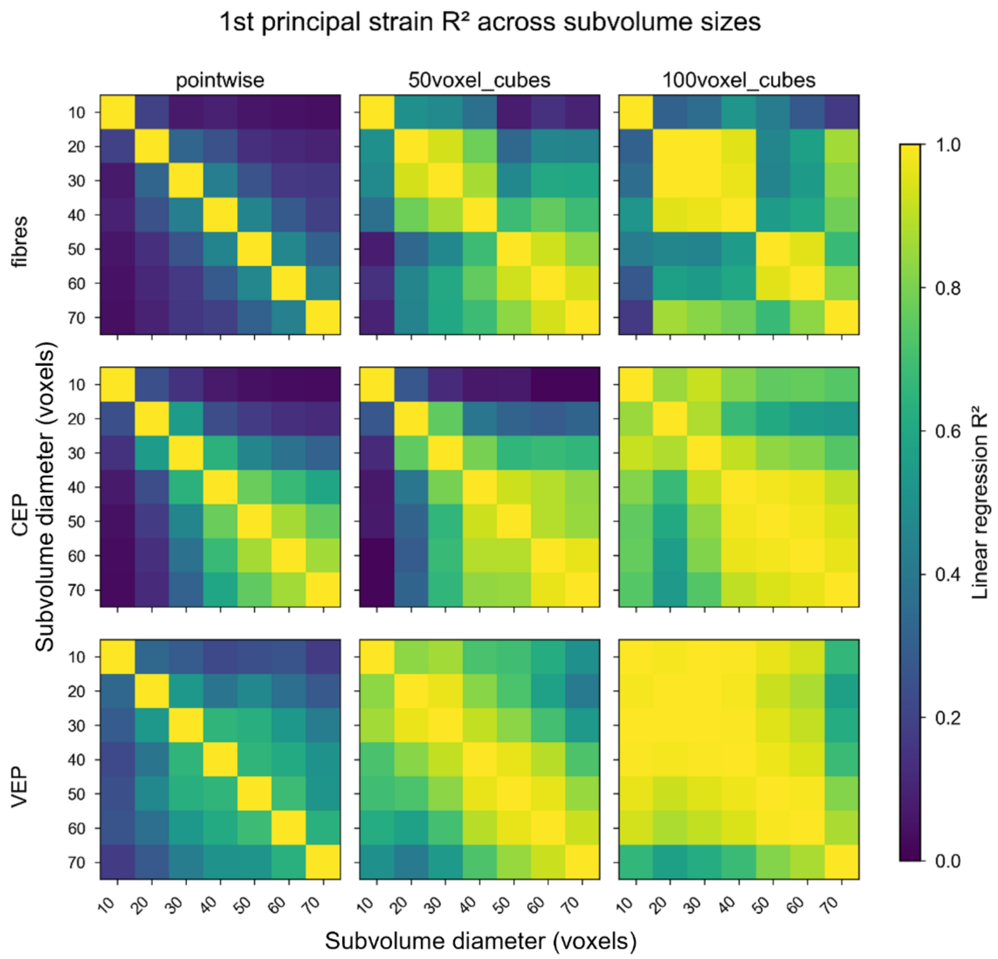


**Supplementary Figure 19 – Linear regression analysis of 1st principal strain fields across different subvolume sizes and spatial averaging length scales for each tissue.**

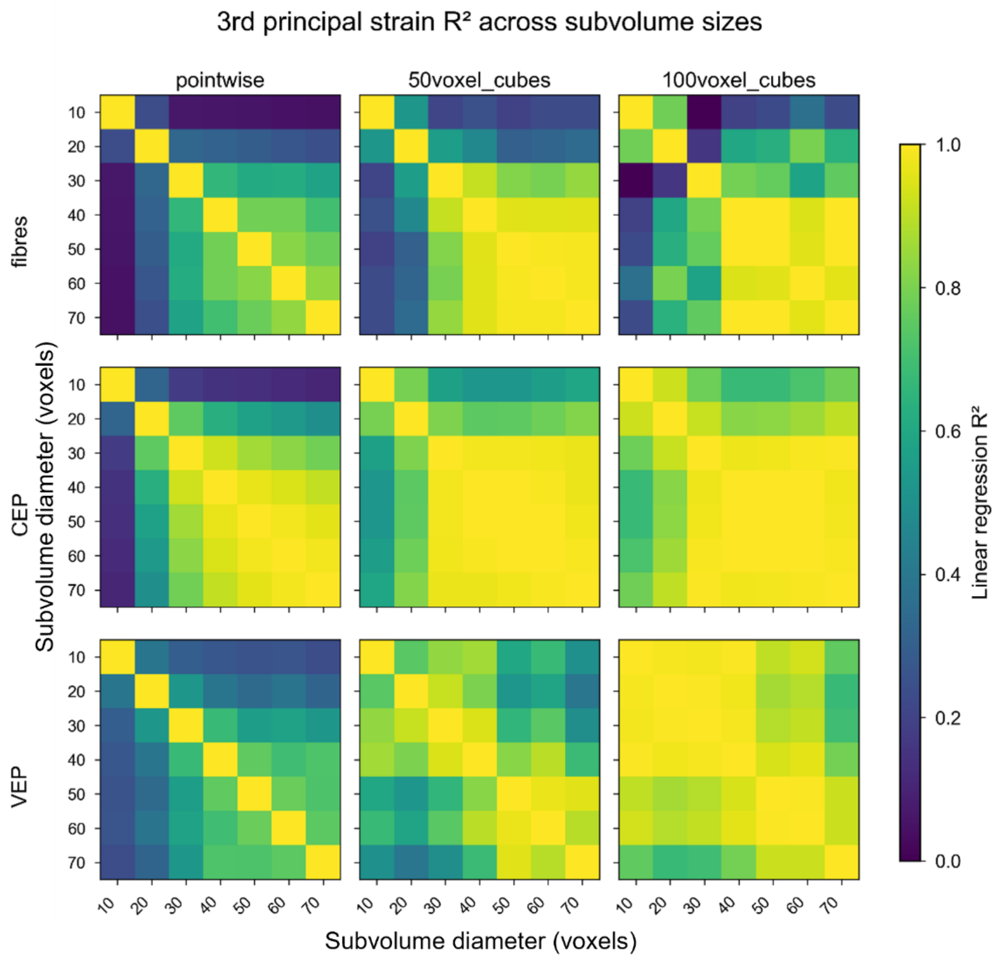


**Supplementary Figure 20 – Linear regression analysis of 3rd principal strain fields across different subvolume sizes and spatial averaging length scales for each tissue.**

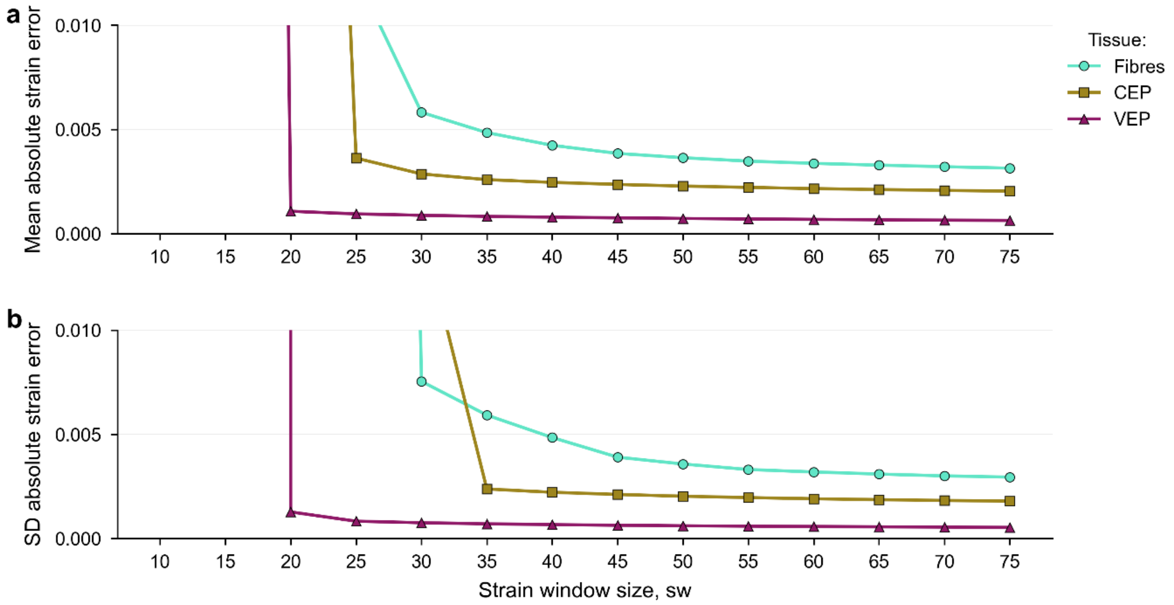


c

| Tissue | Lowest strain MAER | Lowest strain SDER |
|---|---|---|
| Fibres | 0.00314<br>sw 75 | 0.00294<br>sw 75 |
| CEP | 0.00203<br>sw 75 | 0.00179<br>sw 75 |
| VEP | 0.000626<br>sw 75 | 0.000527<br>sw 75 |

**Supplementary Figure 21 – a) Mean absolute strain error and b) standard deviation of absolute strain error with increasing strain window size (number of points included in fitting procedure); c) summary table showing the lowest strain MAER and SDER for each tissue type.**
